# Metastatic Single Tumor Cells Evade NK Cell-mediated Killing by Thrombin-mediated Loss of the Activating Ligand CD155/PVR/Necl-5

**DOI:** 10.1101/2021.01.15.426784

**Authors:** Hiroshi Ichise, Shoko Tsukamoto, Tsuyoshi Hirashima, Yoshinobu Konishi, Choji Oki, Shinya Tsukiji, Satoshi Iwano, Atsushi Miyawaki, Kenta Sumiyama, Kenta Terai, Michiyuki Matsuda

## Abstract

Natural killer (NK) cells lyse invading tumor cells to limit metastatic growth in the lung, but how some cancers evade this host protective mechanism to establish a growing lesion is not known. Here we have combined ultra-sensitive bioluminescence whole body imaging with intravital two-photon microscopy involving genetically-encoded biosensors to examine this question. NK cells eliminated disseminated tumor cells from the lung within 24 hrs of arrival, but not thereafter. Intravital dynamic imaging revealed that a disseminated tumor cell in a pulmonary capillary interacts with an NK cell every 2 hrs on average. In the first 4 hrs after tumor cell arrival, 50% of such encounters lead to tumor cell death but after 24 hrs of arrival, nearly 100% of the interactions result in the survival of the tumor cell. This evasion of NK cell surveillance is mediated by thrombin-dependent loss of cell surface CD155/PVR/Necl-5, a ligand for the NK cell activating receptor DNAM-1. This loss prevents the NK cell signaling needed for effective killing of tumor targets. By quantitatively visualizing the evasion of NK cell surveillance, we have uncovered a molecular mechanism for cancer evasion and provided an explanation for the anti-metastatic effect of anticoagulants.

**SUMMARY:** Intravital functional two-photon microscopy reveals that metastatic tumor cells lodged in pulmonary capillaries acquire resistance to patrolling NK cells. Protease-mediated loss of the activating ligand CD155/PVR/Necl-5 on tumor cells prevents NK cells from ERK activation and tumor cell killing.

## INTRODUCTION

Natural killer (NK) cells are innate lymphoid cells that play critical roles in protecting against the development of tumor metastases (Chiossone et al., 2018). In human patients, a higher number of circulating or tumor-infiltrating NK cells is correlated with better patient outcomes (López-Soto et al., 2017). In immunocompetent mice, selective depletion of NK cells markedly increases the metastatic burden (Diefenbach et al., 2001; Smyth et al., 1999). Most of these previous studies relied on the number of macro-metastatic tumors as a single functional endpoint, preventing insight into the step(s) of the metastatic cascade at which NK cells play the most critical role. Metastasis involves the migration of a tumor cell or tumor cell cluster from the primary cancer site through the blood, lodging of the migrating cells in a micro-vessel, and the transmigration of the cell(s) into the tissue parenchyma, where they may either remain dormant or grow into a larger tumor mass (Gupta and Massagué, 2006). Given this sequence of events, it is unknown where NK cell attack on the malignant cells giving rise to a metastatic lesion takes place. Using a newly developed method of intravascular staining of immune cells (Anderson et al., 2014), it was shown that more than 90% of NK cells in the mouse lung are in the vasculature (Secklehner et al., 2019). In agreement with this finding, the major NK cell subset in the lung is similar to that in the peripheral blood (Hayakawa and Smyth, 2006; Marquardt et al., 2017). Meanwhile, it has also been proposed that NK cells in the normal lung are incompetent or hypofunctional (Marquardt et al., 2017; Robinson et al., 1984). Thus, it remains unclear how pulmonary NK cells are able to prevent tumor cells from colonization in the lung.

Inhibition of platelets and coagulation factors has long been associated with the suppression of lung metastasis (Brown, 1973; Gasic et al., 1968; Pearlstein et al., 1984). At least a part of the pro-metastatic effect of the coagulation cascade is attributed to inhibition of NK cells (Gorelik et al., 1984; Nieswandt et al., 1999; Palumbo et al., 2005; Sadallah et al., 2016). But other mechanisms such as enhanced adhesion of tumor cells (Nierodzik et al., 1992), formation of a favorable intravascular metastatic niche (Lucotti et al., 2019), or TGF-β1-mediated immune evasion (Metelli et al., 2020) have also been proposed. Therefore, a method to untangle the metastatic cascade in vivo is needed to quantitatively assess the effect of anti-coagulants on each step and the possible relationship of this anti-metastatic action to the function of NK cells.

Intravital two-photon (2P) microscopy enables visualization of the interaction of immune cells with other immune cells or their targets (Cyster, 2010; Germain et al., 2012; Liew and Kubes, 2015). For example, the dynamic behaviors of NK cells have been demonstrated in the lymph nodes (Bajénoff et al., 2006; Beuneu et al., 2009; Mingozzi et al., 2016) and the tumor microenvironment (Barry et al., 2018; Deguine et al., 2010). Moreover, the development of genetically-encoded biosensors for signaling molecules has paved the way to monitoring of cellular activation status in not only normal but also pathological tissues (Conway et al., 2017; Terai et al., 2019).

Here, we have combined bioluminescence whole body imaging and intravital 2P microscopy to explore the behavior and functional competence of NK cells in an experimental lung metastasis model. Using an ultra-sensitive bioluminescence system, we followed the fate of intravenously injected tumor cells from 5 min to 10 days. The number of viable disseminated tumor cells in the lung decrease rapidly and reach a nadir within 12**–**24 hrs in an NK cell-dependent manner. Intravital 2P microscopy demonstrates that a static tumor cell in a pulmonary capillary is contacted by a crawling NK cell approximately every 2 hrs. Importantly, the probability of NK cell activation and subsequent elimination of the lodged tumor cell decreases rapidly after 24 hrs of arrival in the lung capillary bed. We show that this evasion of NK cell surveillance is caused by thrombin-dependent shedding of CD155/PVR/Necl-5 (hereafter simply Necl-5), a ligand for the NK cell activating receptor DNAM-1. This loss of surface activating ligand limits the signaling needed to invoke NK cytotoxicity, thus protecting the tumor cell and enabling formation of a growing metastatic lesion. Anti-coagulants promote tumor killing by NK cells by limiting this loss of activating ligand.

## RESULTS

### NK Cells Eliminate Disseminated Tumor Cells Within 12–24 hrs After the Entry into the Lung

Development of an extremely bright bioluminescence imaging system, AkaBLI (Iwano et al., 2018), enabled us to visualize the acute phase of lung metastasis and to explore the role of NK cells in the elimination of disseminated tumor cells. B16F10 melanoma cells were transduced with Akaluc luciferase, and the resulting cells, called B16-Akaluc cells hereafter, were injected intravenously into syngeneic C57BL/6 mice that had previously been injected with either an anti-asialo GM1 (αAGM1 hereafter) antibody or an isotype control antibody. The pretreatment with αAGM1 removed more than 97% of NK cells from the spleen and the lung (Figure S1A and S1B). Immediately after the intravenous injection of tumor cells, we administered AkaLumine luciferin intraperitoneally (i.p.) and started bioluminescence imaging under anesthesia (Figure 1A and Supplementary Video 1). A bioluminescence signal in the control mice was observed almost exclusively in the lung and decreased rapidly in the first 20 min, then gradually thereafter. In the αAGM1-treated mice, the bioluminescence signal dropped rapidly as observed in the control mice; however, the decrease was substantially reduced after 20 min as compared to control animals. Thus, in the initial phase (< 180 min) there are at least two mechanisms that eliminate melanoma cells from the lung. The rapid elimination of melanoma cells (< 20 min) may reflect flushing away by the blood flow or shear-stress-mediated cell death. The slow component of the elimination (> 20 min) observed in the control mice is caused primarily by NK cells.

**Figure 1:**
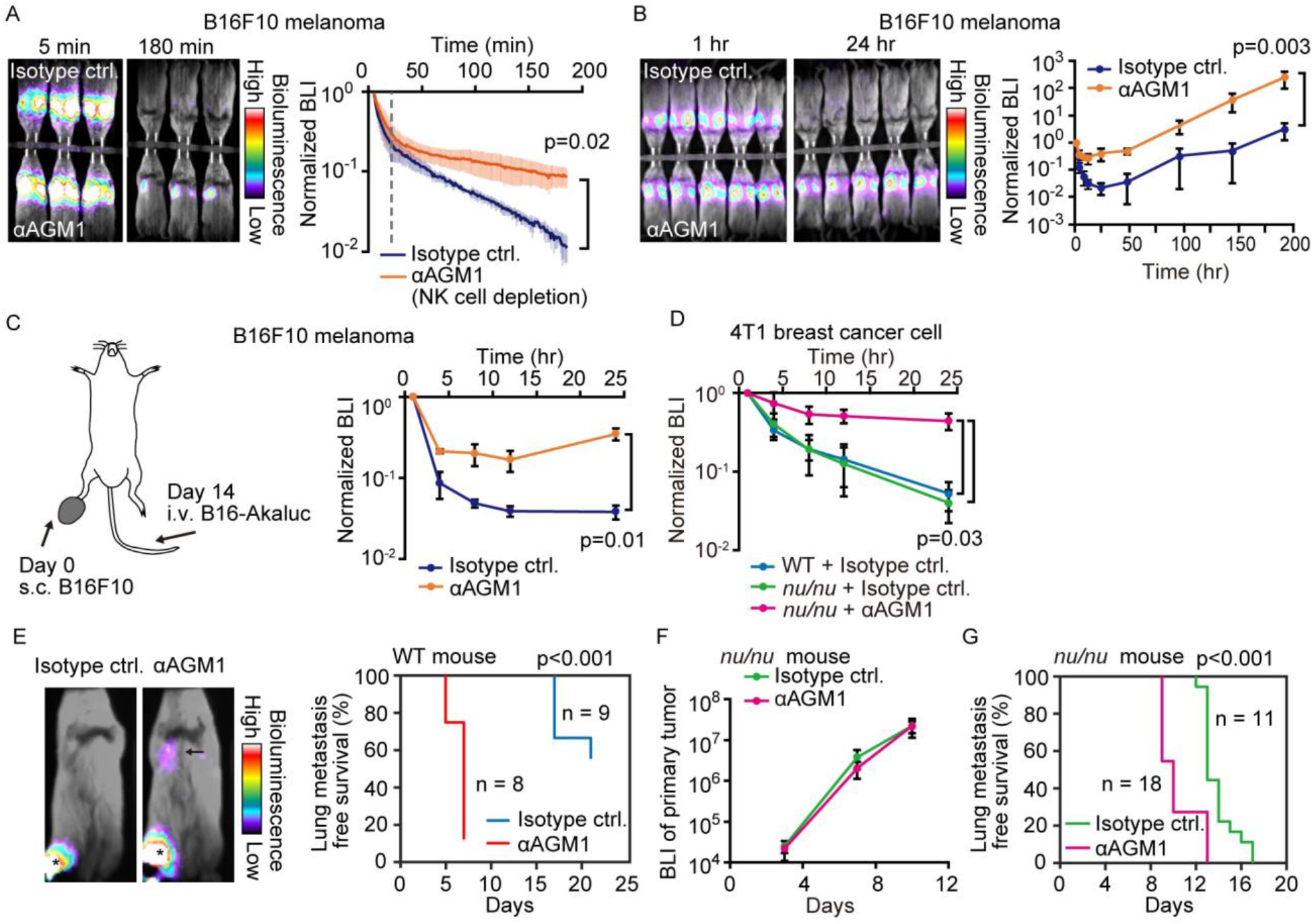
NK Cells Eliminate a Subset of Metastatic Tumor Cells from the Lung Within 24 hrs of Arrival. (a,b) B6 mice were pretreated with either control antibody or αAGM1. Representative merged images of the bright field and the bioluminescence images of mice intravenously injected with 5×10^5^ B16-Akaluc cells are shown. (A) The substrate was i.p. administered immediately after injection of tumor cells. Image acquisition was started at 5 min after tumor injection. See also Supplementary Video 1. Bioluminescence intensity (BLI) is normalized to that at 5 min and plotted over time. Data are representative of 2 independent experiments with 3 mice per group and are shown as means ± SD. A dotted line represents 20 minute. (B) Substrate was administered i.p. before each round of image acquisition. BLI is normalized to that at 1 hr. Data are representative of 3 independent experiments with 4–6 mice per group and shown as means ± SD. (C) B16-Akaluc cells were injected into the tail vein of mice that had been inoculated with B16F10 cells in the foot pad 14 days before. BLI was quantified at the indicated time and normalized to that at 1 hr after injection of the B16-Akaluc cells. Data are representative of 2 independent experiments with 3 mice per group and are represented as means ± SD. (D) BALB/c mice or BALB/c *nu/nu* mice were pretreated with either control antibody or αAGM1. 4T1-Akaluc cells were injected into the tail vein. BLI was quantified at the indicated time and normalized to that at 1 hr after injection of the tumor cells. Data are representative of 2 independent experiments with 4 mice per group and are represented as means ± SD. (E) BALB/c mice were pretreated with either control antibody or αAGM1. Shown are representative merged images of the bright field and the bioluminescence images of mice subcutaneously injected with 5×10^5^ B16-Akaluc cells into footpad. An arrow and asterisks depict a lung metastasis and primary tumors, respectively. Lung metastasis incidence of control antibody- (n = 9) or αAGM1- (n = 8) treated mice. (F) BLI of primary tumor. Data are representative of 2 independent experiments with 3 mice per group and are represented as means ± SD. (G) Identical to (E) except that mice are BALB/c *nu/nu* and the implanted cell number is 1×10^4^.

To explore the NK cell-mediated elimination of tumor cells after the early rapid phase decline (1 hr–8 days), we administered luciferin immediately before each round of imaging (Figure 1B). The bioluminescence signals were normalized to that at 1 hr after B16-Akaluc injection in each mouse. In the control mice, the bioluminescence signal of melanoma cells reached a nadir 24 hrs after tumor cell injection and increased thereafter, indicating proliferation of melanoma cells. On the other hand, in the αAGM1-treated mice, the bioluminescence signal decreased very little after 4 hrs and started increasing after 12 hrs. Importantly, after 24 hrs, we did not observe any significant difference in the relative increase of the bioluminescence signal between the control and αAGM1-treated mice, implying that NK cells eliminate disseminated melanoma cells primarily in the acute phase (< 24 hrs) of lung metastasis. In another model system, B16F10 melanoma cells were inoculated into the foot pad two weeks prior to the intravenous injection of the B16-Akaluc cells (Figure 1C). αAGM1 treatment hampered the rapid decrease of the bioluminescence signal, suggesting that NK cell-mediated elimination of metastatic melanoma cells operates in the melanoma-burdened mice as well.

We extended this approach to other syngeneic mouse tumor cell lines: Braf^V600E^ mutated melanoma cells (Dhomen et al., 2009) and MC-38 colon adenocarcinoma cells (Rosenberg et al., 1986). The rapid decrease of bioluminescence signals was markedly alleviated by αAGM1, supporting the critical role of NK cells in the acute phase (Figure S2A and S2B). It has been reported that the αAGM1 cross-reacts with basophils (Nishikado et al., 2011). Therefore, we repeated the experiment using an αCD200R3 basophil-depleting antibody (Ba103). The basophil depletion did not affect the elimination of melanoma cells (Figure S3A and S3B). Similarly, the roles of circulating monocytes and neutrophils were examined with clodronate liposome and αLy-6G neutrophil-depleting antibody, respectively, and neither the depletion of monocytes nor that of neutrophils mitigated the melanoma elimination within 24 hrs of injection (Figure S3C–S3F).

The effect of T cell immunity on tumor elimination was examined by using BALB/c *nu/nu* mice. The Akaluc luciferase was transduced in BALB/c-derived 4T1 breast cancer cells to generate 4T1-Akaluc cells, which were injected into wild type BALB/c mice or BALB/c *nu/nu* mice (Figure 1D). 4T1-Akaluc cells were eliminated in BALB/c nu/nu mice as efficiently as in wild type mice in an NK cell-dependent manner, indicating that under these conditions, T cell immunity does not contribute to tumor cell reduction. To further explore the NK cell-mediated tumor cell elimination in a spontaneous metastasis model, 4T1-Akaluc cells were inoculated into the foot pad (Figure 1E). The αAGM1 treatment did not affect the growth of the primary tumor, but suppressed lung metastasis (Figure 1F and 1G). These results indicate the critical role of NK cells in the elimination of disseminated tumor cells.

### NK Cell Dynamic Behavior in Pulmonary Capillaries

To better understand how NK cells mediate rapid elimination of metastatic tumor cells we examined their topographic distribution and dynamics in the lung. We first assessed the localization of NK cells by intravascular staining with αCD45 antibody (Anderson et al., 2014), followed by flow cytometry with αCD45, αCD3, and αNKp46 antibodies (Figure S4). Consistent with previous reports (Gasteiger et al., 2015; Secklehner et al., 2019), more than 95% of pulmonary NK cells (CD45^+^, CD3^−^, NKp46^+^) were found in the vasculature, compared to 20% of bone marrow NK cells. To study the dynamics of NK cell-mediated immune surveillance in the lung, we developed a reporter line, NK-tdTomato mice, whose NK cells, or more specifically, NKp46^+^ cells (Narni-Mancinelli et al., 2011), express the fluorescent protein tdTomato. Next, we employed *in vivo* pulmonary imaging by 2P microscopy to observe NK cells *in situ* for up to 12 hrs (Figure 2A). In agreement with the flow cytometric data, most NK cells were found within the capillaries (Figure 2B). NK cells flowing in the capillaries stalled on endothelial cells, crawled a short distance, and jumped back into the flow (Figure 2C and Supplementary Video 2). A histogram of the crawling time exhibited exponential decay with a median of 2.5 min (Figure 2D). NK cells are known to express at least two integrin-family proteins, LFA-1 (LFA-1α/CD18, αLβ2) and Mac-1 (Mac-1/CD18, αMβ2) (Wang et al., 2012). Intravenous injection of the blocking antibody against LFA-1α (αLFA-1α) resulted in a reduction of NK cells on the pulmonary endothelial cells (Figure 2E). To extend this observation, we counted the number of NK cells on the endothelial cells in the presence or absence of αLFA-1α. As expected, αLFA-1α, but not αMac-1, markedly reduced the number of NK cells on the pulmonary endothelial cells (Figure 2F), indicating that the adhesion of NK cells to the pulmonary endothelial cells is mediated primarily by LFA-1.

**Figure 2:**
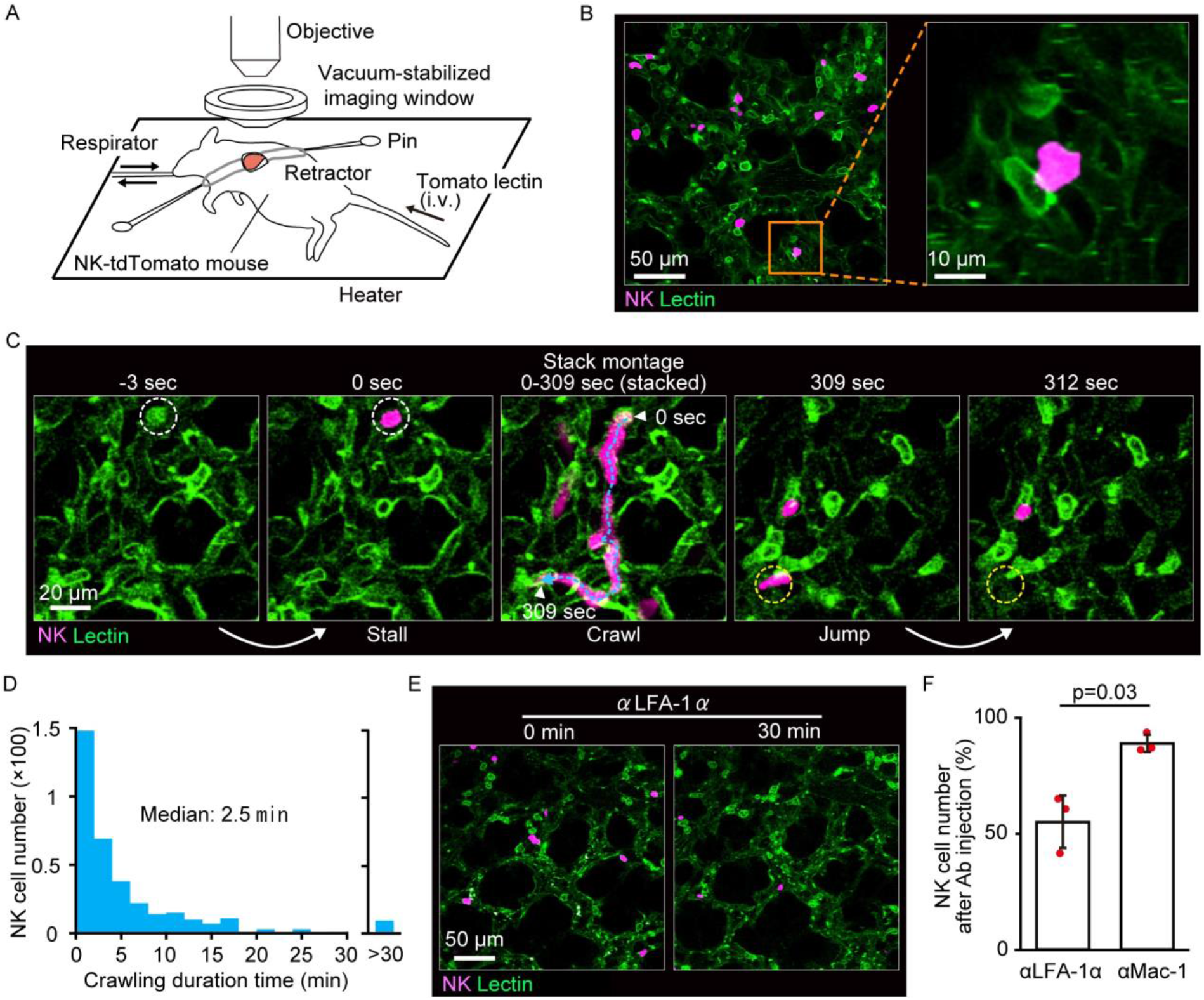
NK Cells Patrol Pulmonary Capillaries in a Stall-Crawl-Jump Manner. (A) A schematic of the intravital imaging system for the lung. The left lobe of the lung was exposed by 5th or 6th intercostal thoracotomy using custom-made retractors and fixed to the objective by a vacuum‐stabilized imaging window. (B) (Left) A micrograph of the lung of an NK-tdTomato mouse, in which NK cells express tdTomato (magenta). Lectin (green) was injected intravenously to stain endothelial cells. (Right) A magnified image of the boxed region in the left panel. (C) A representative time-lapse image of NK cells (magenta). The track of an NK cell is shown with a cyan dotted line and white arrowheads at both ends. White and yellow dotted circles show the positions of stall and jump, respectively. See also Supplementary Video 2. (D) Distribution of the crawling duration times in a 0.25 mm^2^ field of view (FOV). Data are pooled from 2 independent experiments (n=583). (e, f) B6 mice expressing tdTomato in NK cells (magenta) were observed by 2P microscopy. During time-lapse imaging with a 30 sec interval, 100 μg of αLFA-1α or αMac-1 antibody was intravenously injected. The number of NK cells in the 0.25 mm^2^ FOV was counted 0–10 min before and 30 min after antibody injection. The percentage of NK cells after versus before antibody injection is shown in (F). Data were pooled from 3 independent experiments and represented as means ± SD. n=3 mice for each group.

### Intra-pulmonary NK Cell Patrolling and Tumor Interaction Dynamics

To understand better the kinetics of NK cell migration within pulmonary vessels and their interactions with lodged tumor cells, we examined the three-dimensional trajectory of the crawling NK cells in the presence and absence of B16F10 cells (Figure 3A–3C). In this experiment, we used B16-SCAT3 cells, which expressed the Förster resonance energy transfer (FRET)-based caspase-3 biosensor SCAT3 (Takemoto et al., 2003). The ratio of NK cells versus tumor cells in the field of view (FOV) was set approximately 1:1. We compared the mean square displacement (MSD) of NK cells in the presence vs. absence of tumor cells (Figure 3D). In general, the MSD adopts the asymptotic power-law form: *MSD*~*t*^*α*^, where *α* is the degree of diffusive motion. The motion is classified as normal diffusion when *α*=1 but as anomalous diffusion otherwise (Krummel et al., 2016). Curves fitted with the measured data represent *α*<1 regardless of the presence of tumor cells, *α*=0.56 in the presence and *α*=0.53 in the absence, indicating that the migration mode of NK cells can be classified as sub-diffusive. This is probably because the migration of NK cells is limited to the pulmonary vascular structure. MSD analysis also shows that the displacement from the original position is smaller in the presence than in the absence of tumor cells. This observation is consistent with our present finding that the median instantaneous speed of the crawling NK cells was significantly decelerated in the presence of tumor cells, from 7.8 to 4.8 μm/min (Figure 3E). In accordance with this observation, the median duration time of crawling was markedly increased in the presence of tumor cells, from 5 to 30 min (Figure 3F). Notably, these data exclude regions in which the tumor cells would physically block NK movement in the capillary, suggesting that the dissemination of tumor cells causes global activation of pulmonary endothelial cells, and thereby causes slower crawling of NK cells. Taking advantage of these quantitative imaging data, we summarized parameters regarding the dynamics of pulmonary NK cells (Supplementary Table). The parameters on flowing NK cells were deduced from the NK cell count in blood and the flow rate in pulmonary capillaries. In short, a tumor cell lodged in a pulmonary capillary will be contacted by the flowing and crawling NK cells roughly every 10 min and every 2 hrs, respectively.

**Figure 3:**
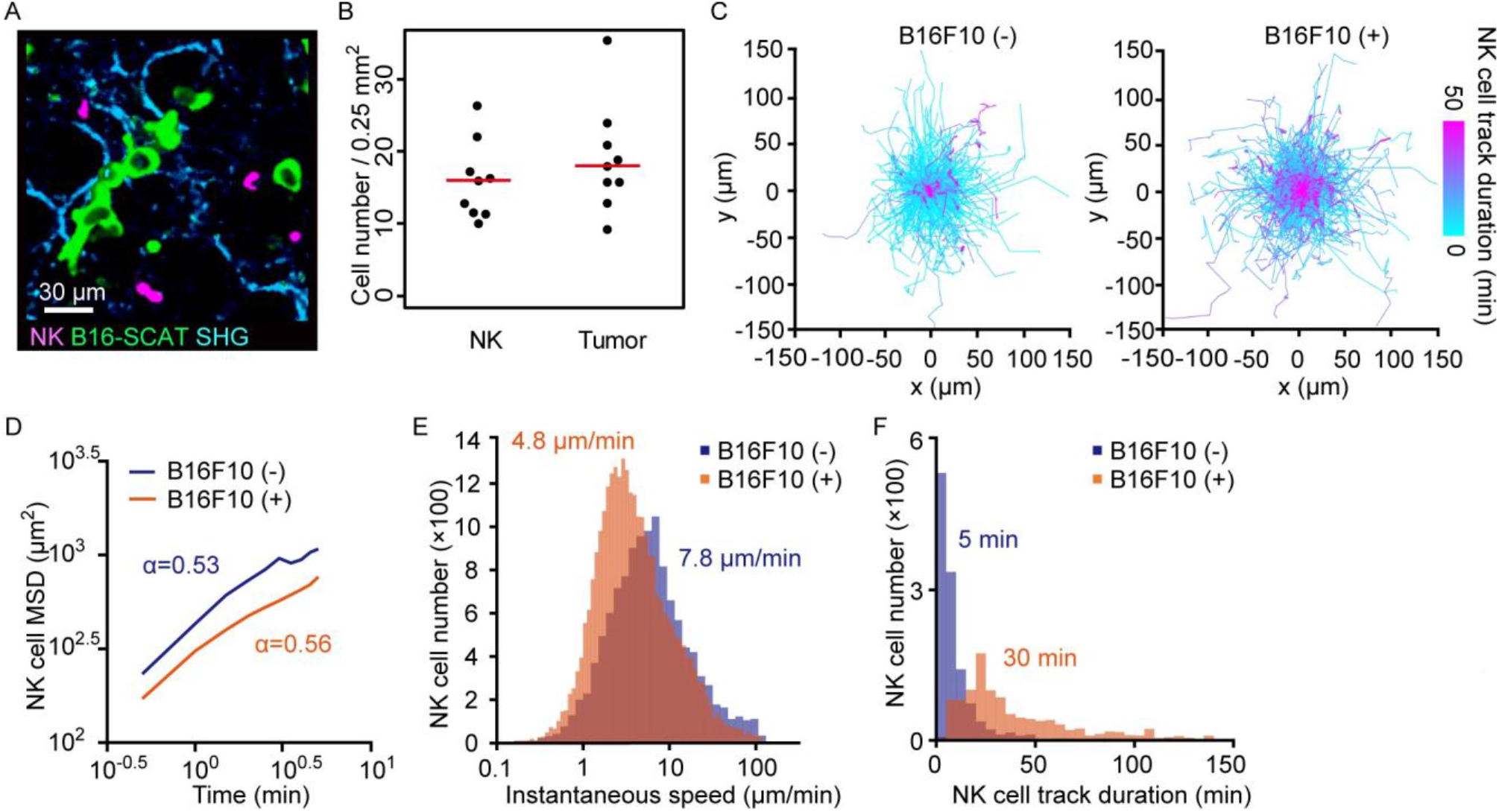
NK Cells Patrol Capillaries Deliberately in the Presence of Melanoma. NK-tdTomato mice were injected with 1.5×10^6^ B16-SCAT3 cells and observed under a 2P microscope for 2 hrs from 6 hrs after injection. NK-tdTomato mice without any treatment were used as the control. (A) A micrograph of the lung of an NK-tdTomato mouse. B16-SCAT3 cells, green; NK cells, magenta. (B) The average number of NK cells and tumor cells in each FOV at 10–35 μm from the pleura 10 min after intravenous injection of B16F10 cells (n=9). The red lines represent the mean. (C) Trajectories of crawling NK cells in the presence (left) or absence (right) of B16F10 cells. For 3D tracking, images of a 0.25 mm^2^ FOV and 25 μm thickness at 10–35 μm from the pleura were acquired every 30 sec for 120 min. Shown here are the trajectories of NK cells projected onto the XY plane. Each track is shown in pseudo-color based on track duration. Data are from 3 independent experiments for each condition. n=1,127 cells in the absence and n=718 cells in the presence of tumor cells. (d–f) Shown are mean squared displacement (MSD) (D), instantaneous speed (E), and track duration (F).

### Necl-5 and Nectin-2 on Tumor Cells Stimulate NK Cell Signaling Leading to Tumor Cell Killing

To gain more insight into NK-mediated tumor killing in the lung, we visualized tumor cell apoptosis with SCAT3. For the typical B16-SCAT3 cells, caspase-3 was activated 16 min after an NK cell came into contact with the tumor cell (Figure 4A and 4B and Supplementary Video 3). The caspase-3 activation was observed in 18% of contact events within a median of 26 min after the contact (Figure 4C and 4D). To examine the possible cause of this limited extent of measurable cell death after contact with an NK cell, we studied Ca^2+^ influx in tumor cells, which is known to herald apoptosis (Keefe et al., 2005), by using two Ca^2+^ sensors, GCaMP6s (Chen et al., 2013) and R-GECO1 (Zhao et al., 2011). In preliminary *in vitro* experiments, the B16F10 cells expressing R-GECO1 (B16-R-GECO) were co-cultured with NK cells that had been activated by IL-2 *in vitro*. Typically, Ca^2+^ influx was observed within a few min after contact (Figure S5A and S5B). A surge of Ca^2+^ influx was observed only in cells that were doomed to die (Figure S5C); 98% of cells that exhibited Ca^2+^ influx died during the imaging (Figure S5D). With these *in vitro* data in hand, we studied Ca^2+^ influx in metastatic melanoma cells *in vivo*. In a typical example (Figure 4E, 4F and Supplementary Video 4), when an NK cell contacted a B16F10 cell expressing GCaMP6s (B16-GCaMP), Ca^2+^ influx was induced within 3 min. The Ca^2+^ influx was observed with a median lag of 6 min (Figure 4G) in 47% of contact events (Figure 4H). These data suggest that the reason why NK cells failed to induce caspase-3 activation in four-fifths of the tumor cells may be due to the limitation of observation period after the delivery of the lethal hit. To connect these data on tumor cell death with NK cell activation, we used B16F10 cells deficient in expression of CD155/Necl-5/PVR and CD112/Nectin-2 and (Necl-5 and Nectin-2, hereafter), the ligands for the activating receptor DNAM-1 on NK cells (Chan et al., 2010). As anticipated, tumor Ca^2+^ influx was almost completely abolished in the *Necl-5* and *Nectin-2-*deficient B16F10 cells (Figure 4H), indicating that damage to tumor cells is dependent on the engagement of DNAM-1 on NK cells.

**Figure 4:**
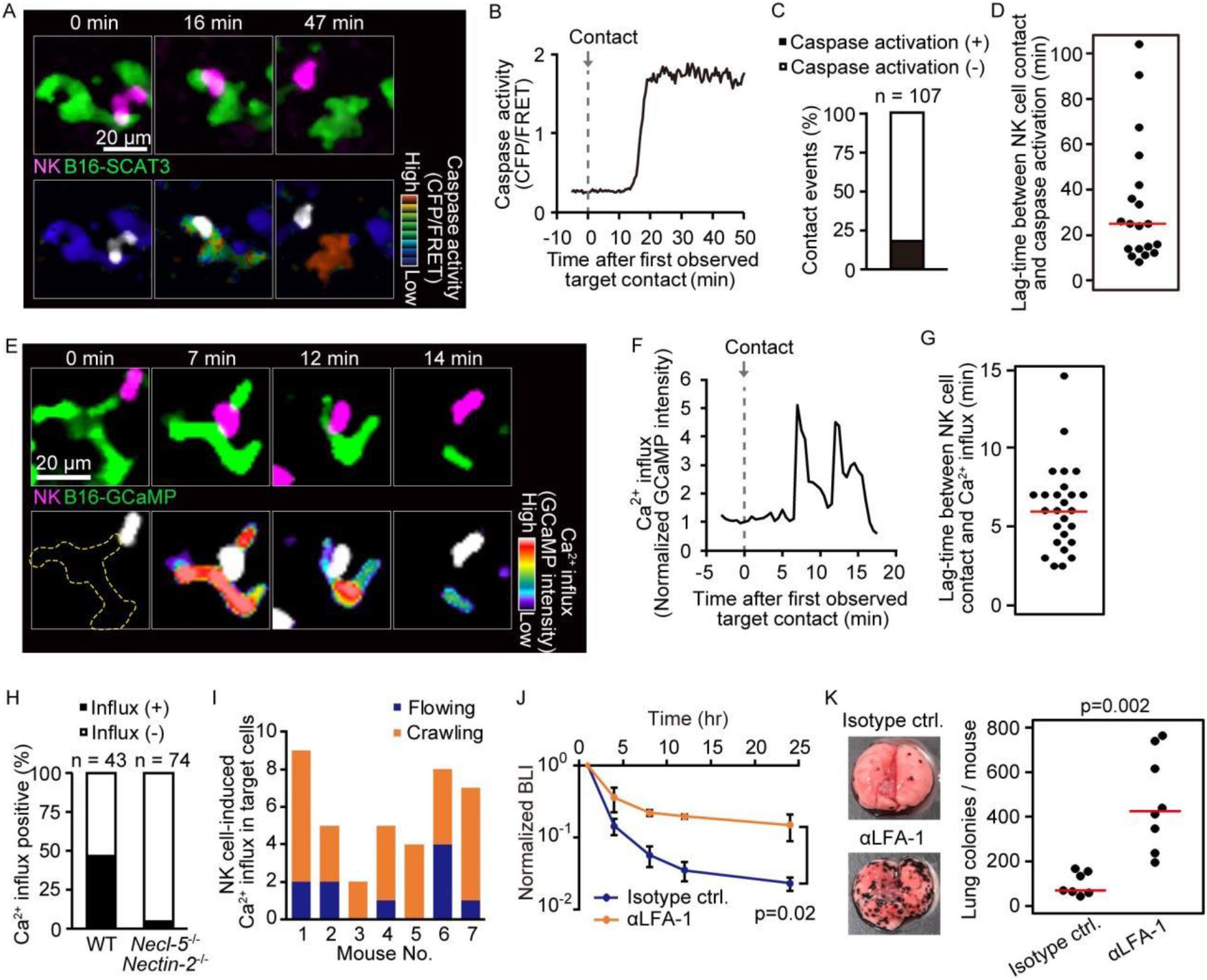
Intravital 2P Imaging with Biosensors Visualizes Apoptosis and Calcium Influx of Tumor Cells Induced by Crawling, but Not Flowing, NK Cells. (A) A representative time-lapse image of a lung of an NK-tdTomato mouse after B16-SCAT3 cell injection. An NK cell and B16-SCAT3 cells are depicted in magenta and green, respectively (top). Bottom, the CFP/FRET ratio in B16-SCAT3 cells is shown in the intensity-modulated display (IMD) mode and an NK cell is shown in white. See also Supplementary Video 3. (B) Quantification of the CFP/FRET ratio in a B16-SCAT3 cell in (A). (C) The percentage of NK cell-tumor cell contacts with or without caspase activation. Data were pooled from 3 independent experiments. (D) Time intervals between NK cell contact and caspase 3 activation in B16-SCAT3 cells. Data are pooled from 3 independent experiments. (E) A representative time-lapse image of a lung of NK-tdTomato mice after B16-GCaMP cell injection. An NK cell and B16-GCaMP cell are depicted in magenta and green, respectively (top). Bottom, GCaMP6s intensity in a B16-GCaMP cell is displayed in pseudo-color and an NK cell is shown in white. See also Supplementary Video 4. (F) Quantification of GCaMP6s intensity shown in (E). (G) Time intervals between NK cell contact and Ca^2+^ influx in B16-GCaMP cells. Data were pooled from 4 independent experiments. Red lines represent the median. (H) Comparison of the number of NK cell contacts that were followed by Ca^2+^ influx between the WT and *Necl-5*^*−/−*^*/Nectin-2*^*−/−*^. Data were pooled from 4 (WT) and 2 (*Necl-5*^*−/−*^*/Nectin-2*^*−/−*^) independent experiments. (I)) In 7 independent experiments, 40 contact events with calcium influx were observed and classified into those caused by crawling or flowing NK cells. (J)) αLFA-1α or isotype control antibody was intravenously administered 2 hrs before injection of 5×10^5^ B16-Akaluc cells. The bioluminescence signals are normalized to those of 1 hr. Data are representative of 2 independent experiments with 3–4 mice per group and presented as means ± SD. (K) Representative macroscopic images of the metastasis to the lung and number of metastatic nodules per lung are shown. Red lines represent the median. Data were pooled from 2 independent experiments. Control, n=7; αLFA-1α, n=8.

With these data in hand we could return to the question of whether flowing or crawling NK cells are responsible for tumor cell death. Crawling NK cells accounted for 77% of Ca^2+^ influx events in the melanoma cells (Figure 4I). To reveal the impact of the crawling NK cells, NK cells αLFA-1α was used to inhibit NK cell attachment to the pulmonary capillaries. This markedly attenuated the melanoma elimination not only within 24 hrs, but also after 10 days (Figure 4J and 4K). Our finding suggests that LFA-1 is required for the crawling of NK cells on the endothelial cells, and, thereby, for the immune surveillance against metastatic tumor cells. B16F10 cells do not express the LFA-1 ligands, ICAM-1 and ICAM-2 (Figure S6), arguing against the possibility that αLFA-1α directly impairs the association of NK cells with B16F10 cells, a conclusion also consistent with a previous study that demonstrated that LFA-1 deficiency in NK cells did not abrogate the *in vitro* killing capacity against B16F10 cells (Zhang et al., 2015).

### Tumor-mediated Stimulation of NK Cells Declines after Several Hours of Pulmonary Residence

To track tumor cell induction of NK cell activation, a key step in the killing process, we took advantage of evidence that engagement of LFA-1 or DNAM-1 with their ligands results in the activation of extracellular signal-regulated kinase (ERK) (Perez et al., 2003; Zhang et al., 2015). We isolated NK cells from transgenic mice expressing a FRET-based ERK biosensor (Komatsu et al., 2018), and co-cultured them with B16F10 cells *in vitro*. Quantification of ERK activity in each NK cell during contact with the tumor cells demonstrated a positive correlation between ERK activation in NK cells and apoptosis in tumor cells (Figure S7A). Apoptosis was induced in 47% of tumor cells that were in contact with the ERK-activated NK cells (Figure S7B). On the other hand, apoptosis was never observed in the tumor cells in contact with the NK cells that failed to exhibit ERK activation. In agreement with this observation, an inhibitor for MAPK/ERK kinase PD0325901, called MEKi hereafter, suppressed NK cell-mediated apoptosis (Figure S7C). When NK cells were sorted into DNAM-1^+^ and DNAM-1^−^, ERK was activated more potently in DNAM-1^+^ NK cells than in DNAM-1^−^ NK cells (Figure S7D), supporting the critical role of DNAM-1 in ERK activation.

The link between ERK activation and elimination of metastatic tumor cells was examined in vivo using bioluminescence imaging. At 1 hr before and 8 hrs after the injection of B16-Akaluc cells, MEKi or solvent was administered i.p. into mice. MEKi significantly attenuated the rapid decrease of the injected tumor cell number within 24 hrs (Figure 5A) and the number of lung colonies of MEKi-treated mice was significantly greater than in control mice (Figure 5B). At the same time, we observed that MEKi had no additive effect in αAGM1-treated mice in the early (< 24 hrs) time courses (Figure 5A). Although interpretation is limited by the action of the soluble inhibitor on cells other than NK cells, these additional data are consistent with our imaging data and the idea that ERK-dependent NK cell activation contributes to the elimination of disseminated tumor cells.

**Figure 5:**
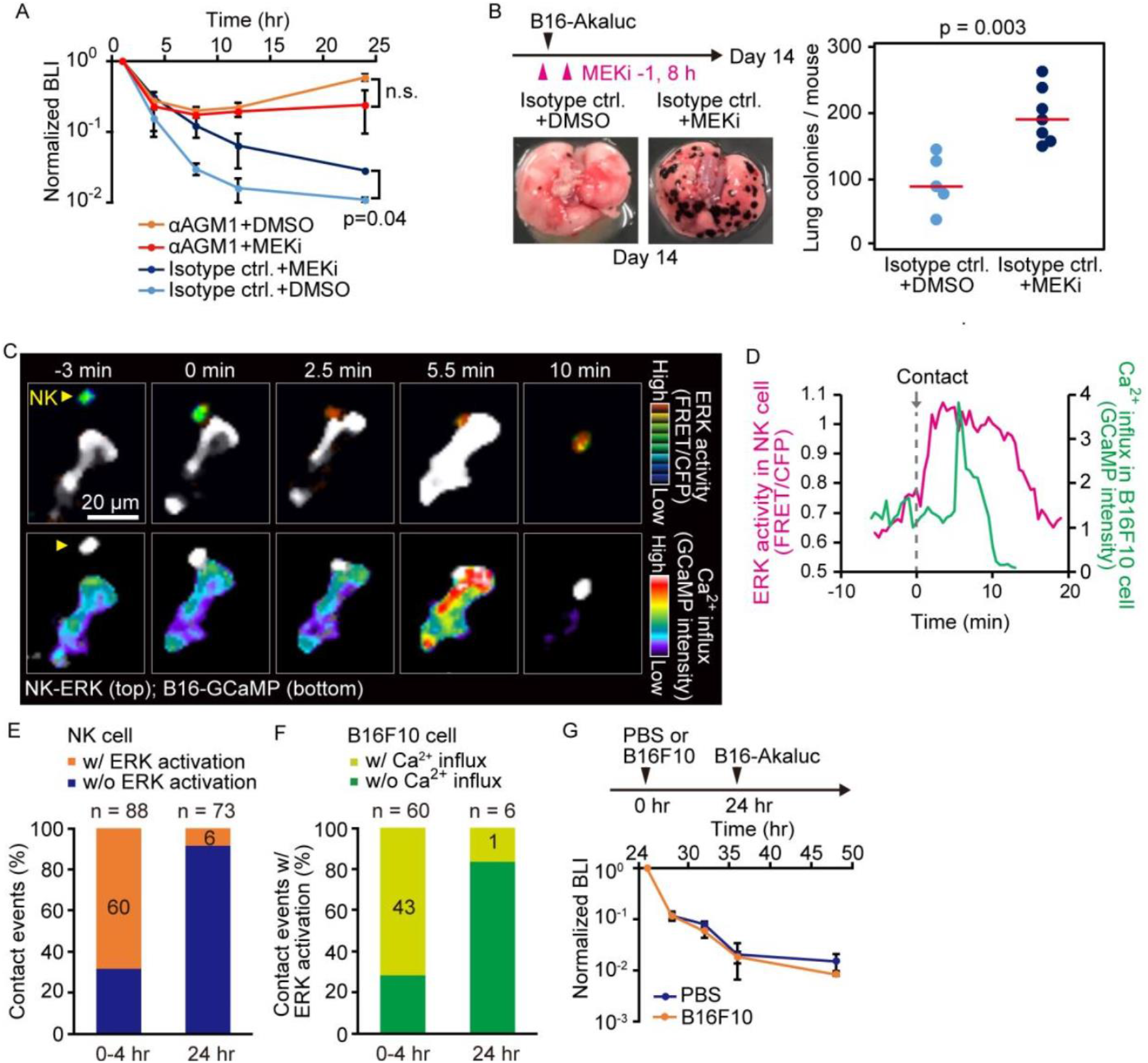
Contact-Induced ERK Activation in NK Cells Is a Necessary Event in Induction of Apoptosis in Tumor Cells in the First 4 Hrs, but Not after 24 Hrs. (A) B6 mice were pretreated with either control antibody or αAGM1 and intravenously injected with 5×10^5^ B16-Akaluc cells. At 1 hr before and 8 hrs after injection, an MEK inhibitor (MEKi) or DMSO was administered i.p. Time courses of the signals, which are normalized to those at 1 hr after tumor injection for each mouse. Data are representative of 2 independent experiments and shown as means ± SD. n.s., not significant. (B) Macroscopic images were acquired at day 14. The number of metastatic colonies are shown. Control, n=5; MEKi, n=7. (C) A time-lapse image of the lung of an NK-ERK mouse expressing the FRET biosensor for ERK. The mouse was intravenously injected with B16-GCaMP cells. Top, FRET/CFP ratio of an NK cell (yellow arrowhead) is shown in IMD mode. A B16-GCaMP cell is shown in white. Bottom, GCaMP6s intensity is displayed in pseudo-color. The NK cell is shown in white. (D) Time course of the FRET/CFP ratio in the NK cell and CaMP6s intensity in the B16-GCaMP cell. (E) Activation probability of ERK in the NK cells upon target cell contact at 0-4 hrs or 24 hrs after tumor injection. Data were pooled from 3 independent experiments. (F) The probability of NK cells that exhibited ERK activation with or without induction of Ca^2+^ influx in the target tumor cells at 0-4 hrs or 24 hrs after tumor injection. Data were pooled from 3 independent experiments. (G) B16-Akaluc cells were injected into the tail vein of mice that had been injected PBS or B16F10 cells into tail vein 24 hrs before. BLI was quantified at the indicated time and normalized to that at 1 hr after injection of B16-Akaluc cells. Data are representative of 2 independent experiments with 3 mice per group and are represented as means ± SD.

With these data in hand, we next proceeded to visualize ERK activation *in vivo*. For this, we developed reporter mice whose NK cells express the FRET biosensor for ERK, hereinafter called NK-ERK mice. Intravenous injection of B16-GCaMP cells into NK-ERK mice allowed for simultaneous observation of ERK activity in NK cells and Ca^2+^ influx in melanoma cells. In a representative example, ERK was activated 2.5 min after an NK cell’s contact with a melanoma cell (Figure 5C, upper panel; Figure 5D, magenta), followed by Ca^2+^ influx in the melanoma cells at 5.5 min and cell death at 10 min (Figure 5C, lower panel; Figure 5D, green). ERK activation, defined by a more than 30% increase in the FRET/CFP ratio, was observed within 3 min in 60 NK cells during 88 contact events (Figure S8A; Figure 5E). Ca^2+^ influx was observed at a median of 4 min in 43 of the 60 tumor cells that came into contact with the NK cell having ERK activation (Figure S8B; Figure 5F). We did not observe Ca^2+^ influx in 28 tumor cells that were touched by the NK cells that failed to show evidence of ERK activation. This result supports the notion that ERK activation in NK cells contributes to the induction of apoptosis in the target tumor cells. Importantly, 24 hrs after injection, the probability of ERK activation and Ca^2+^ influx was markedly decreased, indicating NK cells lose the capacity to activate in response to capillary-lodged tumor cells tumor cells (Figure 5C and 5D).

To understand the basis for this loss of NK activity by 24 hrs after tumor arrival in the lung, B16F10 melanoma cells without Akaluc were injected 24 hrs before the injection of B16-Akaluc cells (Figure 5G). We did not observe any effect of the pre-injected B16F10 cells on the time course of clearance of the B16-Akaluc cells, indicating that NK cells do not lose tumoricidal activity, but that over time, the tumor cells acquire the capacity to evade NK cell surveillance.

### Thrombin-Mediated Shedding of Necl-5 Causes Evasion of NK Cell Surveillance

DNAM-1-mediated signaling is indispensable for tumoricidal activity of NK cells, leading us to reason that cell surface expression of the DNAM-1 ligands Necl-5 and Nectin-2 may diminish over this time period. Because the expression of Nectin-2 was significantly less than that of Necl-5 in B16F10 melanoma cells, we focused on Necl-5. As anticipated, cell surface expression of Necl-5 was markedly decreased in tumor cells isolated from the lungs 24 hrs after injection (Figure 6A, 6B). To explore the basis for this change, we expressed a recombinant Necl-5 fused extracellularly to the mScarlet fluorescent protein and intracellularly to the mNeonGreen fluorescent protein (Figure 6C). The ratio of extracellular mScarlet versus intracellular mNeonGreen was markedly reduced in 24 hrs, indicating loss of extracellular domain, possibly by cleavage or shedding (Figure 6D, 6E). A clue to the mechanism of this Necl-5 loss came from evidence that inhibition of serine proteases in the coagulation cascade have anti-tumor effects in rodent models and human patients (Francisco and Palumbo, 2019; Nierodzik and Karpatkin, 2006). Warfarin, an anti-vitamin K drug potently accelerated the acute elimination of tumor cells (Figure 6F). Similarly, specific inhibitors of factor Xa and thrombin, edoxaban and dabigatran etexilate, respectively, also promoted the elimination of tumor cells (Fig 6G, 6H). Notably, dabigatran etexilate did not have significant effect in αAGM1-treated mice (Figure 6H), indicating that the anti-metastatic effect is mediated by NK cells. In agreement with these functional results, thrombin cleaved off the extracellular domain of Necl-5 from tumor cells *in vitro* (Figure 6I). Several mechanisms have been proposed for coagulation cascade-mediated NK cell inhibition (Francisco and Palumbo, 2019), including that platelet-tumor aggregates physically protect tumor cells from NK cells (Cluxton et al., 2019; Nierodzik and Karpatkin, 2006). However, we rarely observed microthrombi around the disseminated tumor cells, even when a microthrombus was easily induced by laser ablation in the vicinity of the tumor cell (Figure S9 and Supplementary Video 5). Considering the role of platelets as the platform for coagulation factors, platelets may play a pro-metastatic role by promoting thrombin activation and thereby Necl-5 shedding from the metastatic tumor cells.

**Figure 6:**
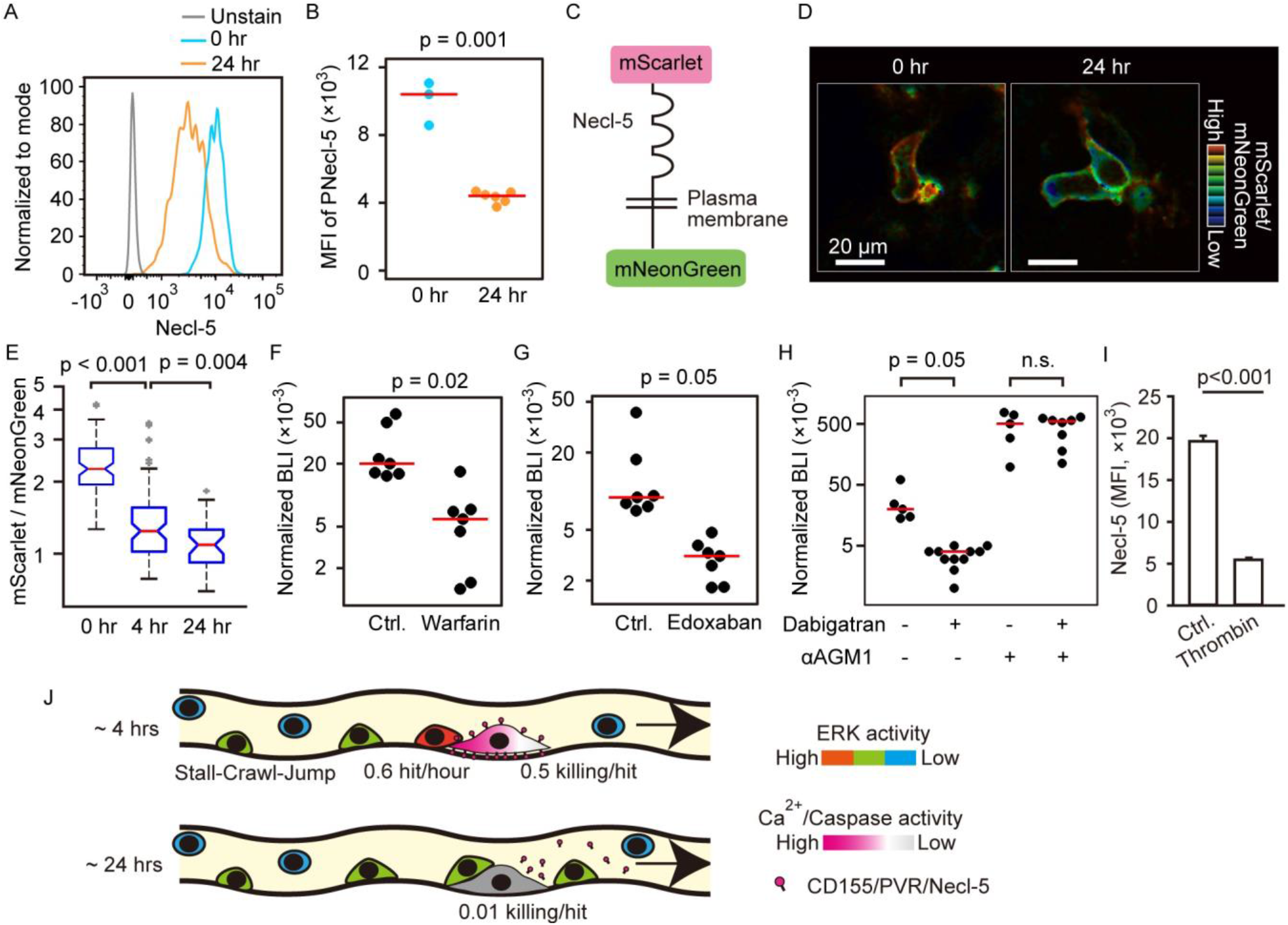
Shedding of Necl-5 Causes Evasion of NK Cell Surveillance. (a, b) B16-Akaluc cells were injected into the tail vein and the expression level of Necl-5 on survived tumor cells was analyzed at 24 hrs after dissemination. The MFI of Necl-5 in tumor cells injected 0 hr or 24 hrs before is shown in (B). Red lines represent the median. Data were pooled from 2 independent experiments. (C) Schematic representation of the Necl-5-ScNeo fusion protein. (D) The representative images of mScarlet/mNeonGreen ratio in the B16F10 cells expressing Necl-5-ScNeo at 0.5 hrs and 24 hrs after injection are shown in the IMD mode. The quantified mScarlet/mNeonGreen ratio in the transmembrane in indicated time point is shown in (E). Mice were treated in their drinking water with 5 mg/L warfarin at least for 5 days and intravenously injected with 5×10^5^ B16-Akaluc cells. The BLI at 24 hrs, which normalized to those at 1 hr after tumor injection for each mouse are shown. Red lines represent the median. Data were pooled from two independent experiments. (g, h) Mice were intravenously injected with 5×10^5^ B16-Akaluc cells. At 1 hr before and 12 hrs after tumor injection, edoxaban, dabigatran or vehicle was orally administered to mice. For NK cell depletion, mice were pretreated with either control antibody or αAGM1. The BLI at 24 hrs, which normalized to those at 1 hr after tumor injection for each mouse are shown. Red lines represent the median. Data were pooled from two independent experiments. (I) Flow cytometric analysis of B16F10 cells treated with recombinant thrombin for 3 hrs. Data are representative of 2 independent experiments with 3 wells per group and are represented as means ± SD. (J) Evasion of NK cell surveillance by shedding of Necl-5.

## DISCUSSION

Despite the established role of NK cells in the prevention of metastasis (López-Soto et al., 2017), the step(s) of the metastatic cascade at which NK cells eliminate disseminated tumor cells remain unknown. Here, we adopted the AkaBLI system, by which even a single Akaluc-expressing tumor cell could be detected in mice (Iwano et al., 2018), and followed the fate of intravenously injected tumor cells from 5 min to 10 days after the injection of tumor cells (Figure 1). In agreement with previous studies (Grundy et al., 2007; Hinuma et al., 1987), the number of tumor cells decreased rapidly in an NK cell-dependent manner. However, we noticed that as early as 24 hrs after injection, tumor cells started to increase and formed macrometastatic nodules, irrespective of the presence or absence of NK cells. Our results demonstrate that the principal role of NK cells in the prevention of lung metastasis is to destroy tumor cells shortly after their arrival and lodging in the pulmonary vasculature, but not thereafter. Using intravital imaging, Headley et al. found that tumor microparticles are rapidly ingested by myeloid cells to evoke an immune response (Headley et al., 2016). However, depletion of myeloid lineage cells did not affect the development of metastasis in the models we employed (Figure S3), excluding a role of the myeloid cells in the clearance of disseminated tumor cells in the immediate-early phase. Patrolling behavior similar to that of NK cells has also been observed for intravascular monocytes in the skin (Auffray et al., 2007). Like NK cells, the crawling monocytes require LFA-1 for their crawling. However, again, depletion of monocytes did not prevent the rapid clearance of disseminated tumor cells (Figure S3). Thus, the rapid elimination of the disseminated tumor cells within 24 hrs is dependent primarily on NK cells in our models.

The crawling NK cells induce pre-apoptotic calcium influx in approximately 50% of the tumor cells that they contact (Figure 4H), indicating that pulmonary NK cells can eliminate the disseminated tumor cells as efficiently as do chimeric antigen receptor (CAR) T cells (Cazaux et al., 2019). Our findings are superficially inconsistent with reports that lung NK cells exhibit highly differentiated and hypofunctional phenotypes *in vitro* (Hayakawa and Smyth, 2006; Marquardt et al., 2017; Robinson et al., 1984). This discrepancy probably arises from the fact that most of the tumor cells were killed by NK cells crawling on the pulmonary endothelial cells, but not by those flowing in the blood. NK cells adhere to the endothelial cells in an LFA-1-dependent manner (Figure 4J). Binding of LFA-1 to its ligand ICAM-1 induces reorganization of the actin cytoskeleton and polarization of NK cells, which is a prerequisite for subsequent redistribution of cytotoxic granules toward the bound targets (Mace et al., 2009). It is likely that LFA-1 engagement of ICAM on pulmonary endothelial cells contributes to the pre-activation status of NK cells. In this regard, the slow migration speed of NK cells in the presence of tumor cells may indicate the increased fraction of active NK cells (Figure 3).

In the B16F10 melanoma metastasis model, DNAM-1 expression in NK cells is crucial for the rejection of tumor cells (Gilfillan et al., 2008). In agreement with a previous report (Zhang et al., 2015), we have shown that, upon target cell engagement, ERK is rapidly activated in DNAM-1^+^ NK cells, but not DNAM-1^−^ NK cells (Figure S7D). Simultaneous *in vivo* visualization of ERK activity in NK cells and Ca^2+^ influx in tumor cells revealed highly efficient tumor cell killing by NK cells showing evidence of effective signaling based on ERK activation (Figure 5E, 5F). Only about 60% of lung NK cells express DNAM-1 (Tahara-Hanaoka et al., 2005). However, most of the tumor cells were killed by the NK cells crawling on the pulmonary endothelial cells, but not by those flowing in the blood. Because DNAM-1 serves as an adhesion molecule (Kim et al., 2017; Shibuya et al., 1996), the crawling NK cells may be biased to DNAM-1^+^ NK cells, consistent with the potent cytotoxic activity of the crawling cells.

Shedding of ligands for the activating receptors of NK cells has been documented for NKG2D (Raulet et al., 2013). The NKG2D ligands, MICA, MICB, and ULBP2 are cleaved by matrix metalloproteases (MMP), leading to evasion of NK surveillance. It is also reported that platelets can promote the MMP-mediated shedding of the NKG2D ligands in vitro (Maurer et al., 2018). However, to the best of our knowledge, involvement of thrombin or factor Xa in the shedding of the ligands for activating receptors such as Necl-5 has not been reported. Interestingly, Necl-5 contributes to cell adhesion as does DNAM-1 (Takai et al., 2008). Therefore, Necl-5 on tumor cells is a double-edge sword, which facilitates adhesion to the lung capillary but also invokes NK cell attack. To evade the NK cell surveillance, after anchoring to the capillaries, tumor cells are able to take advantage of the capacity of thrombin to strip Necl-5 from the cell surface within 24 hrs after adhesion (Figure 6J). This new understanding of how NK cell surveillance of tumor cells in the micro-circulation can be evaded provides a rationale for the development of new ways to inhibit growth of clinically significant lung metastases.

## MATERIALS AND METHODS

### Plasmids

Plasmids encoding R-GECO1, SCAT3, GCaMP6s, mScarlet were obtained from Takeharu Nagai (Zhao et al., 2011), Masayuki Miura (Takemoto et al., 2003), and Addgene (plasmid # 40753 for GCaMP6s; #85044 for mScarlet; Cambridge, MA), respectively. The cDNAs of mNeonGreen (Shaner et al., 2013) was synthesized with codon optimization by GeneArt (Thermo Fisher Scientific, Waltham, MA). A plasmid encoding tdTomato was obtained from Takara Bio (#632533; Kusatsu, Japan). pCSIIhyg-R-GECO1 and pCSIIbsr-GCaMP6s, lentiviral vectors for R-GECO1 and GCaMP6s, respectively, were constructed by inserting cDNAs into pCSII-based lentiviral vectors (Miyoshi et al., 1998) with IRES-hyg (hygromycin B-resistance gene) or IRES-bsr (blasticidin S-resistance gene). psPAX2 (Addgene Plasmid #12260) and pCMV-VSV-G-RSV-Rev (provided by Hiroyuki Miyoshi at RIKEN) were used for the lentivirus production. To generate pPBbsr-SCAT3-NES, cDNA coding the SCAT3 fused with the nuclear export signal (NES) (LQLPPLERLTLD) of the HIV-1 rev protein (Fischer et al., 1995) was subcloned into pPBbsr, a PiggyBac transposon vector with IRES-bsr (Yusa et al., 2009). pPBbsr2-Venus-Akaluc was described previously (Iwano et al., 2018). To generate pPBbsr2-Necl-5-ScNeo, cDNA coding the signal peptide of Necl-5 (a.a. 1-28), mScarlet, Necl-5 (a.a. 29-408), and mNeonGreen were PCR-amplified and assembled into pPBbsr2 vector by using In-Fusion system (Takara Bio USA, Inc., Mountain View, CA). pCMV-mPBase (obtained from the Wellcome Trust Sanger Institute) was co-transfected with pPB vector to establish stable cell lines. To generate pT2Aneo-tdTomato-CAAX, cDNA encoding tdTomato fused with the CAAX domain of the KRas protein (a.a. 170–189) was subcloned into pT2Aneo vector (obtained from Koichi Kawakami (Kawakami et al., 2004)), a Tol2 transposon vector with IRES-neo (neomycin-resistance gene). For the gene knockout, lentiCRISPR v2 vector (Addgene, #52961) was used. To generate transgenic mice, pT2ADW-lox-mCherry-hyBRET-ERK-NLS was constructed by assembling cDNAs of hyBRET-ERK-NLS (Komatsu et al., 2018), mCherry, and the NES sequence of the HIV-1 rev protein into pT2ADW vector (Komatsu et al., 2018) by In-Fusion cloning (Takara Bio).

### Reagents

PD0325901 (FUJIFILM Wako Pure Chemical Corporation, Osaka, Japan) was applied as an MEK inhibitor. Warfarin, Lixianar (edoxaban), and Prazaxa (dabigatran etexilate) was obtained from Eisai Co., Ltd. (Tokyo, Japan), DAIICHI SANKYO COMPANY, LIMITED (Tokyo, Japan), and Boehringer Ingelheim GmbH (Ingelheim, Germany), and used as a vitamin K inhibitor, factor Xa (FXa) inhibitor, and thrombin inhibitor, respectively. AkaLumine-HCl (TokeOni) was obtained from Kurogane Kasei Co., Ltd. (Nagoya, Japan) or synthesized as previously described (Kuchimaru et al., 2016) and used as the substrate of Akaluc. Collagenase type IV and DNase I were obtained from Worthington Biochemicals (Lakewood, NJ) and Roche (Basel, Switzerland), respectively. A LIVE/DEAD Fixable Red Dead Cell Stain Kit (Thermo Fisher Scientific) or 7-AAD (BD Bioscience) was used to stain dead cells in flow cytometry. DyLight 488-labeled Lycopersicon esculentum lectin was purchased from Vector Laboratories (Burlingame, CA). Recombinant mouse protein C and factor Xa were obtained from R&D Systems, Inc. (Minneapolis, MN). Recombinant mouse thrombin was obtained from antibodies-online GmbH (Aachen, Germany).

### Antibodies

The following antibodies were used for staining: BV510 or FITC anti-CD45 (30-F11), APC-Cy7 anti-CD3 (145-2C11), PerCP-Cy5.5 anti-NK1.1 (PK136), APC or PE-Cy7 anti-DNAM-1 (10E5), PE anti-F4/80 (BM8), APC anti-CD11b (M1/70), PE anti-NKp46 (29A1.4), PE anti-Ly-6G (1A8), PE anti-c-Kit (2B8), APC anti-CD49b (DX5), PE-Cy7 anti-CD200R3 (Ba13), PE anti-ICAM-1 (YN1/1.7.4), Alexa 488 anti-ICAM-2 (3C4), PE-anti PVR/Necl-5 (TX56), PE-anti-human-Nectin-2 (TX31) (all from BioLegend, San Diego, CA). The following antibodies were used for *in vivo* blocking experiments: anti-LFA-1α (M17/4) and anti-Mac-1 (M1/70) (both from Bio X Cell, West Lebanon, NH). The following antibodies or reagents were used for *in vivo* cell depletion: anti-asialo GM1 (FujiFilm Wako Pure Chemical Corporation, Osaka, Japan), Rabbit IgG isotype control (Thermo Fisher Scientific), anti-Ly6G (1A8; BioLegend), Rat IgG2a isotype control (BioLegend), anti-CD200R3 (Ba103; Hycult Biotech, Uden, Nederland), Rat IgG2b isotype control (BioLegend), clodronate liposomes (Hygieia Bioscience, Osaka, Japan) or control liposome (Hygieia Bioscience).

### Flow cytometry analysis

After staining, cells suspended in PBS containing 3% FBS were analyzed and/or sorted with a FACS Aria IIu cell sorter (Becton Dickinson, Franklin Lakes, NJ). The following combinations of lasers and emission filters were used for the detection of fluorescence: for the fluorescence of BV510, a 405 nm laser and a DF530/30 filter (Omega Optical); for the fluorescence of FITC and Alexa 488, a 488 nm laser and a DF530/30 filter; for the fluorescence of PerCP/Cy5.5 and 7-AAD, a 488 nm laser and a DF695/40 filter (Omega Optical); for the fluorescence of PE, a 561 nm laser and a DF582/15 filter (Omega Optical); for the LIVE/DEAD Fixable Red Dead Cell Stain Kit, a 561 nm laser and a DF610/20 filter (Omega Optical); for the fluorescence of PE-Cy7, a 561 nm laser and a DF780/60 filter (Omega Optical); for the fluorescence of APC, a 633 nm laser and a DF660/20 filter (Omega Optical); and for the fluorescence of APC-Cy7, a 633 nm laser and a DF780/60 filter. Cells were first gated for size and granularity to exclude cell debris and aggregates, and dead cells were excluded by LIVE/DEAD Fixable Red Dead Cell Stain Kit or 7-AAD. Data analysis was performed using FlowJo software (Tree Star, Ashland, OR).

### Staining of intravascular NK cells of the bone marrow and lung

Mice were subjected to intravenous injection of the 3 μg αCD45 antibody 3 min before sacrifice. Bone marrow cells were harvested from the femur by flushing the bone with 5 ml RPMI containing 10% FBS, 100 units/ml penicillin, and 100 μg/ml streptomycin. Red blood cells were removed by lysis with ACK lysing buffer and centrifugation at 500g for 5 min at 4°**C**. A single-cell suspension of the lung cells was generated by mincing the resected lungs with scissors and incubating the minced tissue in RPMI containing 200 U/ml collagenase type IV and 5 U/ml DNase I for 30 min at 37 ºC. The lysed tissue was then passed through a ϕ40 μm cell strainer. The flow-through fraction was washed with PBS by centrifugation at 500g for 5 min at 4°**C**. Cells were analyzed by flow cytometry as described above.

### Tumor cells

The melanoma cell line B16F10 was purchased from the Cell Resource Center for Biomedical Research (Sendai, Japan). The MC-38 mouse colon adenocarcinoma cell line was provided by Takeshi Setoyama and Tsutomu Chiba at Kyoto University. The BRAF^V600E^ melanoma cell line (Dhomen et al., 2009) was provided by Caetano Reis e Sousa at the Francis Crick Institute. 4T1 mammary tumor cells were purchased from ATCC (Manassas, VA) and maintained on a collagen-coated dish (AGC Techno Glass, Tokyo, Japan). All cell lines were cultured in complete RPMI medium (Thermo Fisher Scientific) containing 10% FBS (Sigma-Aldrich, St. Louis, MO), 1 mM sodium pyruvate (Thermo Fisher Scientific), 50 μM 2-mercaptoethanol (Nacalai Tesque), 1% GlutaMAX solution (Thermo Fisher Scientific), 1% MEM Non-Essential Amino Acids (Thermo Fisher Scientific), 10 mM HEPES solution (Thermo Fisher Scientific), 50 μM 2-mercaptoethanol (Nacalai Tesque), 100 units/ml penicillin, and 100 μg/ml streptomycin (Nacalai Tesque, Kyoto, Japan). Mycoplasma contamination is regularly checked using PlasmoTest mycoplasma detection kit (InvivoGen, San Diego, CA).

### Establishment of stable cell lines

To prepare the lentivirus, pCSIIhyg-R-GECO1 or pCSIIbsr-GCaMP6s was cotransfected with psPAX2 and pCMV-VSV-G-RSV-Rev into Lenti-X 293T cells (Clontech, Mountain View, CA) with Polyethylenimine “Max” (Mw 40,000; Polysciences, Warrington, PA). Virus-containing media were harvested 48 hrs after transfection, filtered, and used to infect B16F10 cells to yield B16-R-GECO cells and B16-GCaMP cells. For the transposon-mediated gene transfer, pPBbsr-SCAT3-NES, pPBbsr2-Venus-Akaluc, or pPBbsr2-Necl-5-ScNeo was cotransfected with pCMV-mPBase into B16F10 cells by using Lipofectamin 3000 reagent (Thermo Fisher Scientific), yielding B16-SCAT3 cells, B16-Akaluc cells, and B16-Necl-5-ScNeo cells, respectively. pT2Aneo-tdTomato-CAAX was cotransfeted with pCS-TP into B16-GCaMP6 cells by using Lipofectamin 3000 reagent, yielding B16-GCaMP-tdTomato-CAAX cells. pPBbsr2-Venus-Akaluc was cotransfected with pCMV-mPBase into BRAF^V600E^ melanoma cells, MC-38 cells, and 4T1 cells by using Lipofectamin 3000 reagent. Cells were selected with either 10 μg/ml blasticidin S (Calbiochem, San Diego, CA) or 100 μg/ml hygromycin B (FujiFilm Wako Pure Chemical Corporation).

### CRISPR/Cas9-mediated establishment of KO cell lines

For CRISPR/Cas9-mediated KO of *tyrosinase* (*Tyr*), *Necl-5*, and nectin cell adhesion molecule 2 *(Nectin-2)*, single guide RNAs (sgRNA) targeting the first or second exon were designed using the CRISPRdirect program (http://crispr.dbcls.jp/). For the establishment of double knockout cells, a puromycin-resistant gene in lentiCRISPR v2 vector was replaced with a bleomycin-resistant gene. The targeting sequences were as follows: *Tyr*, GGGTGGATGACCGTGAGTCC; *Necl-5*, GCTGGTGCCCTACAATTCGAC; *Nectin-2*, GACTGCGGCCCGGGCCATGGG. Annealed oligo DNAs for the sgRNAs were cloned into the lentiCRISPR v2 vector. The sgRNA/Cas9 cassettes were introduced into cells by lentiviral gene transfer. Infected cells were selected by 3.0 μg/ml puromycin (InvivoGen) or 100 μg/ml zeocin (Thermo Fisher Scientific). Cells deficient for *Necl-5* and *Nectin-2* were sorted by a FACS Aria IIu cell sorter and used without single cell cloning. *Tyr*-KO cells were subjected to single cell cloning and examined for the frame-shift mutation by nucleotide sequencing.

### Mice

C57BL/6N (hereinafter called B6) mice, BALB/c mice, and BALB/c *nu/nu* (hereinafter called nude) mice were purchased from Shimizu Laboratory Supplies (Kyoto, Japan) and bred at the Institute of Laboratory Animals, Graduate School of Medicine, Kyoto University under specific-pathogen-free conditions. B6N-Tyrc-Brd/BrdCrCrl (hereinafter called B6 Albino) mice were obtained from Charles River Laboratories. Mice were used at the age of 6–18 weeks.

B6.Cg-Gt(ROSA)26Sortm9(CAG-tdTomato)Hze/J mice (JAX 007909) were obtained from the Jackson Laboratory (Bar Harbor, ME). B6(Cg)-Ncr1tm1.1(icre)Viv/Orl mice (Narni-Mancinelli et al., 2011) (hereinafter called *NKp46*^iCre^ mice) were obtained from INFRAFRONTIER (Oberschleissheim, Germany). Transgenic mice expressing hyBRET-ERK-NLS have been described previously (Komatsu et al., 2018). B6.Cg-Gt(ROSA)26Sortm9(CAG-tdTomato)Hze/J mice were crossed with *NKp46*^iCre^ mice for NK cell-specific expression of tdTomato, resulting in *NKp46*^iCre^/ B6.Cg-Gt(ROSA)26Sortm9(CAG-tdTomato)Hze/J mice (hereinafter called NK-tdTomato mice). Tg(lox-mCherry-hyBRET-ERK-NLS) mice were crossed with *NKp46*^iCre^ mice for NK cell-specific expression of hyBRET-ERK-NLS, resulting in *NKp46*^iCre^/ Tg(lox-mCherry-hyBRET-ERK-NLS) mice (hereinafter called NK-ERK mice). The animal protocols were reviewed and approved by the Animal Care and Use Committee of Kyoto University Graduate School of Medicine (approval no. 19090).

### Generation of transgenic mice

Transgenic mice were generated by Tol2-mediated gene transfer as previously described (Sumiyama et al., 2010). Briefly, fertilized eggs derived from B6 mice were microinjected with a mixture of Tol2 transposase mRNA and pT2ADW-lox-mCherry-hyBRET-ERK plasmid. The offspring mice, named Tg(lox-mCherry-hyBRET-ERK-NLS), were then backcrossed with B6 Albino mice for at least 3 generations. Newborn mice were illuminated with a green LED and inspected for red fluorescence through a red filter LED530-3WRF (Optocode, Tokyo).

### In vivo cell depletion

To deplete NK cells, mice were injected i.p. with 20 μg αAGM1 or rabbit IgG isotype control antibody on 1 and/or 2 days before tumor cell injection. In some experiments, the antibody administration was repeated on days 0 and 7 after tumor cell injection. To deplete basophils, 30 μg αCD200R3 (Ba103) or rat IgG2b isotype control antibody was injected i.p. at 1 day before tumor cell injection. To deplete neutrophils, 200 μg anti-Ly6G (1A8) or rat IgG2a isotype control was injected i.p. at 1 day before tumor cell injection. To deplete monocytes and macrophages, clodronate liposome (50 mg/kg) or control liposome was injected i.p. at 1 day before tumor cell injection. The efficiency of the depletion was assessed by flow cytometry.

### In vivo blocking of integrins

NK-tdTomato mice were injected i.v. with 100 μg of αLFA-1α or αMac-1 antibody during image acquisition. To visualize the vascular structure, 50 μg of DyLight 488-labeled lectin was intravenously injected. The number of NK cells in the 0.25 mm^2^ FOV was counted 0–10 min before and 30 min after antibody injection. Image acquisition and analysis were carried out with MetaMorph software (Molecular Devices LLC, Sunnyvale, CA).

### Bioluminescence imaging of lung metastasis

B6 albino, BALB/c, or Nude mice (female, 8-10 weeks old) were used in this experiment. Akaluc-expressing cells were suspended in PBS at 2×10^6^ cells/ml, and 250 µl of the suspension was injected intravenously 60 min before imaging. Mice were anesthetized with 1.5% isoflurane (FujiFilm Wako Pure Chemical Corporation) inhalation and placed on a custom-made heating plate in the supine position. Immediately after the administration (i.p.) of 100 μl of 5 mM AkaLumine-HCl, bioluminescent images were acquired using a MIIS system (Molecular Devices Japan, Tokyo) equipped with an iXon Ultra EMCCD camera (Oxford Instruments, Belfast, UK) and a lens (MDJ-G25F095, φ16 mm, F: 0.95; Tokyo Parts Center, Saitama, Japan). Images were acquired under the following condition: binning, 4; EM gain, 1,000. During the interval of long-term observation (more than 1 hr), mice were recovered from anesthesia and, immediately before observation, anesthetized again and administered AkaLumine as described. In some experiments, 4×10^5^ B16F10 melanoma cells in 50 μl PBS containing 50% Geltrex (Thermo Fisher Scientific) were injected into the foot pad of 6 weeks old female mice. After two weeks, B16-Akaluc cells were injected from tail vein. For serial injection, 5×10^5^ B16-Akaluc cells were injected into the tail vein of mice that had been injected PBS or B16F10 cells into tail vein 24 hrs before. For blocking of LFA-1, mice were injected with 100 μg of αLFA-1α antibody or an isotype control antibody, rat IgG2a, 2 hrs before tumor injection. For MEK inhibition, mice were injected with PD0325901 i.p., 1 hr before and 8 hrs after tumor injection. For vitamin K-dependent protease inhibition, mice were treated in their drinking water with 2.5–5 mg/L warfarin dissolved in an ethanol solution. Control mice were treated with the ethanol vehicle solution only. Solutions were prepared fresh every 3–4days. The treatment started at least day −5 before tumour injection and continued until the end point of the experiment. For factor X or thrombin inhibition, mice were orally administrated with 20 mg/kg edoxaban or 330 mg/kg dabigatran etexilate twice a day. Prothrombine time was measured on blood samples collected from warfarin, edoxaban, and dabigatran etexilate-treated mice before starting experiment using CoaguChek XS system (Roche). For continuous observation of less than 3 hrs, 100 μl of 15 mM AkaLumine-HCl was administered to the mice immediately after tumor injection. Acquisition of bioluminescent images was started at 5 min after tumor injection and repeated every 1 min. Image acquisition and analysis were carried out with MetaMorph software.

### Bioluminescence imaging of spontaneous lung metastasis

BALB/c or Nude mice were inoculated with 5×10^5^ or 1×10^4^ 4T1-Akaluc cells suspended in 50 μl PBS containing 50% Geltrex at the footpad of right hind limb, respectively. Every 2 to 3 days, bioluminescence images were acquired immediately after the administration (i.p.) of 100 μl of 5 mM AkaLumine-HCl. To mask bioluminescence signals from the primary tumor site, right hind limb was covered with black silicon clay. Images were acquired under the following condition: binning, 4; no EM gain, Exposure: 180 sec. Image acquisition and analysis were carried out with MetaMorph software.

### Counting of macroscopic lung metastasis

Single-cell suspensions of B16-Akaluc cells (5×10^5^) were injected intravenously into mice. The lungs were harvested on day 14 or 15, and tumor nodules were counted under a dissection microscope.

### Intravital pulmonary imaging by two‐photon excitation microscopy

Lung intravital imaging was performed as described previously (Kamioka et al., 2017) with some modifications. Mice were anesthetized by 1.5% isoflurane inhalation and placed in the right lateral position on an electric heating pad. The body temperature was maintained at 36°C using a heating pad with a BWT-100A rectal thermometer feedback controller (Bio Research Center, Nagoya, Japan). The mice were anesthetized with 1.0% isoflurane supplied through a tracheostomy tube Surflo indwelling catheter 22G (Terumo, Tokyo, Japan) connected to an MK‐V100 artificial respirator (Muromachi Kikai, Tokyo, Japan). The respirator condition was as follows: O_2_ and air gas ratio, 80:20; beats per min, 55; gas flow, 35 ml/min; inspiratory/expiratory ratio, 3:2. The left lung lobe was exposed by 5th or 6th intercostal thoracotomy with custom-made retractors. A custom‐made vacuum‐stabilized imaging window was placed over the lung. Minimal suction (0.3–0.4 bar) was applied to stabilize the lung against the coverslip. Mice were observed with an FV1200MPE‐ BX61WI upright two-photon excitation microscope (Olympus, Tokyo, Japan) equipped with an XLPLN 25XW‐MP 25X/1.05 water‐immersion objective lens (Olympus) and an InSight DeepSee laser. Image areas of 500×500 μm to a depth of 25 μm were acquired every 30 sec for 60–120 min, with Z steps at 2.5 μm. Images of 512×512 pixels were scanned at 2 μs/pixel with 1.0–1.2× digital zoom. The excitation wavelengths for cyan fluorescent protein, green fluorescent protein and tdTomato were 840, 930, and 1040 nm, respectively. We used an IR‐cut filter, BA685RIF‐3 (Olympus), two dichroic mirrors, DM505 and DM570 (Olympus), and four emission filters, BA460‐ 500 (Olympus) for cyan fluorescent protein, BA495‐540 (Olympus) for green fluorescent protein, BA520‐560 (Olympus) for yellow fluorescent protein, and 645/60 (Chroma Technology Corp., Bellows Falls, VT) for tdTomato fluorescence. All movies were median-filtered for noise reduction. Image analysis was carried out with Imaris (Bitplane, Belfast, UK) and MetaMorph software.

### Visualization of signaling molecule activity in vivo

To detect caspase activity and Ca^2+^ influx in tumor cells under a 2P microscope, 1.5–2.0×10^6^ B16-SCAT3 cells, B16-GCaMP cells, or B16-GCaMP-tdTomato-CAAX cells with *Tyr* deficiency were administered to NK-tdTomato mice through the tail vein. After tumor injection, a 500×500 μm field of view (FOV) at a depth of 25 μm was imaged every 30 sec for 4 to 6 hrs. For simultaneous observation of ERK activity in NK cells and Ca^2+^ influx in melanoma cells, 1.5–2.0×10^6^ B16-GCaMP cells were administered to NK-ERK mice through the tail veil. After tumor injection, a 500×500 μm FOV at a depth of 25 μm was imaged every 30 sec for 6 hrs. The excitation wavelength for CFP and GCaMP was 840 nm. Fluorescent images were acquired with three channels using the following filters and mirrors: an infrared (IR)-cut filter, BA685RIF-3, two dichroic mirrors, DM505 and DM570, and four emission filters, FF01-425/30 (Semrock, Rochester, NY) for the second harmonic generation channel (SHG Ch), BA460-500 for the CFP Ch, BA520-560 for the FRET and GCaMP6s Ch. For the characterization of flowing and crawling NK cells, 1.5-2×10^6^ B16-SCAT3 cells or B16-GCaMP cells were administered to NK-tdTomato mice through the tail veil. From after 2 hrs, images of a 0.25 mm^2^ FOV at a depth of 25 μm were acquired every 30 sec for 2 hrs. Crawling NK cells were defined as cells whose trajectory was recorded in more than 4 frames, i.e., 2 min, before contact with a melanoma cell. Flowing NK cells were defined as cells that were already in contact with the melanoma cells when they first appeared in the FOV. Cells were counted manually by using MetaMorph software.

### Tracking and Motion analysis of NK cells in the lung

For 3D tracking, time-lapse image areas of 500×500 μm and 25 μm thickness at a depth of 10–35 μm were acquired every 30 sec. In some experiments, 1.5–2×10^6^ B16-SCAT3 cells were intravenously injected and images were acquired for 2 hrs after 4 hrs after tumor injection. Image analysis was carried out with Imaris and MetaMorph software. Tracking of NK cells or tumor cells was performed by the 3D tracking function of Imaris. The time-series data of the coordinates were used to calculate track duration, length, speed and displacement. The parameters of 3D tracking by Imaris were as follows: max distance, 30 μm; max frame gap, 2. We used the 3D position of traced cells for the motion analysis. The instantaneous speed *v* was calculated as the speed between two consecutive time frames, i.e., *v* = |**r**(*t*) − **r**(*t* − Δ*t*)|/Δ*t*, where **r** is the position of cells, t is the elapsed time, and Δ*t* is the time interval. We chose Δ*t*=0.5 min. For Figure S4d–f, we ensembled all data over the cells and the elapsed time up to 50 min. The mean square displacement (MSD) of NK cells was calculated by the following equation: *MSD* = 〈|**r**_*i*_(*t*) − **r**_*i*_(0)|^2^〉_*i*_, where **r**_*i*_ is the position of cell *i*, and 〈 〉 represents the average over cells. For the curve fitting in the MSD analysis, we used the nonlinear least-squares solver “lsqcurvefit”, a built-in function of MATLAB (Mathworks Inc., Natick, MA) to determine the exponent parameter of the diffusivity. An NK cell hit on a tumor is defined by the event when an NK cell comes within 10 μm of a tumor cell. The hit probability is obtained by dividing the total number of hit events by the sum of the observation period of each tumor cell. MATLAB scripts and the datasets are available upon request.

### Observation of thrombus in pulmonary capillaries

1.5–2×10^6^ B16-GCaMP-tdTomato-CAAX cells were administered to hyBRET-ERK-NES mice through the tail veil and images were acquired after 24 hrs. During image acquisition, injuries on endothelial walls were generated by momentarily exposing a small area of the vessel wall to a laser of 70 mW power at 840 nm for up to one second. Image analysis was carried out with MetaMorph software.

### In vitro culture of NK cells

NK cells were purified by negative selection from mice splenocytes with an NK cell isolation kit II (Miltenyi Biotec, Bergisch Gladbach, Germany) in accordance with the manufacturer’s instructions. The post-sort purity of NK cells (NK1.1^+^CD3^−^) was >95%. Purified NK cells were plated in 96-well U-bottomed plates (Thermo Fisher Scientific) in complete RMPI medium supplemented with the recombinant murine 1,000 U/ml IL-2 (PeproTech, Rocky Hill, NJ) and cultured for 5 days. In some experiments, DNAM-1^+^ or DNAM-1^−^ NK cells were purified by a FACS Aria Ⅱu on day 2, and further cultured for 3 days. The purity of each NK cell fraction was >98%.

### Time-lapse imaging of in vitro killing of tumor cells

B16-R-GECO cells with *Tyr* deficiency (2×10^4^) were plated on a collagen-coated 96-well glass-base plate (AGC, Tokyo, Japan) and cultured more than 6 hrs to facilitate cell adhesion. Immediately after starting imaging with an epifluorescence microscope, 2×10^4^ NK cells derived from hyBRET-ERK-NLS mice were added to the wells containing adherent target cells. The cells were imaged with an IX81 inverted microscope (Olympus) equipped with a UPlanSApo 40x/0.95 objective lens (Olympus), a PRIME scientific CMOS camera (Photometrics, Tucson, AZ), a Spectra-X light engine (Lumencor, Beaverton, OR), an IX2-ZDC laser-based autofocusing system (Olympus), a MAC5000 controller for filter wheels and XY stage (Ludl Electronic Products, Hawthorne, NY), and an incubation chamber (Tokai Hit, Fujinomiya, Japan). The filters and dichroic mirrors used for time-lapse imaging were as follows: for FRET imaging, an 430/24 (Olympus) excitation filter, an XF2034 (455DRLP) (Omega Optical, Brattleboro, VT) dichroic mirror, and FF01-483/32 (Semrock) and 535/30 (Olympus) for CFP and FRET, respectively. For Red fluorescent protein imaging, 572/35 (Olympus) excitation filters, 89006 (Chroma Technology Corp.) and FF408/504/581/667/762-Di01 (Semrock) dichroic mirrors, and 632/60 (Olympus) emission filters, respectively. MetaMorph software was used for background noise subtraction and image analysis. Background intensities were determined by using an empty culture dish with the same amount of media. After background subtraction, the FRET/CFP ratio images were represented in the intensity-modulated display (IMD) mode. In the IMD mode, eight colors from red to blue were used to represent the FRET/CFP ratio, with the intensity of each color indicating the mean intensity of FRET and CFP channels. To track NK cells, the CFP images were analyzed by using the Fiji TrackMate plugin. From the x and y coordinates, the fluorescence intensity of each cell was isolated and the FRET/CFP ratio was calculated by MATLAB.

### Flow cytometric analysis of disseminated tumor cells in the lungs

1.5–2.0×10^6^ B16-Akaluc cells were intravenously injected and single cell suspension of the lung cells was generated after 24 hrs as describe above. The expression level of Necl-5 was analyzed by FACS Aria IIu cell sorter. Data analysis was performed using FlowJo software.

### Quantification of shedding of extracellular domain of Necl-5 in the lung

1.5–2.0×10^6^ B16-Necl-5-ScNeo cells were intravenously injected and images were acquired after 24 hrs by intravital pulmonary imaging with a two‐photon excitation microscope as describe above. After taking images at 24 hrs, moving to the other field of view without B16-Necl-5-ScNeo cells and 1.0×10^6^ B16-Necl-5-ScNeo cells were newly injected from tail vein. Images of newly injected B16-Necl-5-ScNeo cells were acquired with the same condition with the image acquisition of 24 hrs. The intensity of mScarlet and mNeonGreen in the plasma membrane was isolated by using MetaMorph software and the mScarlet/mNeonGreen ratio was calculated.

### In vitro protease digestion

1.0×10^5^ B16F10 cells or 293T cells were resuspended in serum free RPMI and incubated for 3 hrs at 37°**C** with 100 ug/ml thrombin. The expression level of murine Necl-5 was analyzed by FACS Aria IIu cell sorter. Data analysis was performed using FlowJo software.

### Quantification and statistical analysis

The statistical differences between the two experimental groups were assessed by Welch’s t-test. Kaplan-Meier survival analyses were performed using MATLAB, and the log rank test was used to determine significance.

## ACKNOWLEDGMENTS

We thank J. Miyazaki, T. Nagai (Osaka University), and M. Miura (University of Tokyo) for the plasmids, T. Setoyama, T. Chiba (Kyoto University), and C. Reis e Sousa (Francis Crick Institute) for the cell lines, M. Yanagida (Kyoto University) for the mice, F. Gochi (Kyoto University) for the intercostal thoracotomy, and R. N. Germain (NIAID) and K. Ikuta (Kyoto University) for the critical reading of this manuscript. We are grateful to the members of the Matsuda Laboratory for their helpful input, and to K. Hirano, K. Takakura, A. Kawagishi and Y. Takeshita for their technical assistance. This work was supported by the Kyoto University Live Imaging Center. Financial support was provided by JSPS KAKENHI grant nos. 18K15317 (to H.I.), 15H05949 (to M.M.), 16H06280 (to M.M.), and 19H00993 (to M.M.), AMED grant no. 19gm5010003h0003 (to M.M.), Fugaku Trust for Medical Research (to M.M.), and JST CREST grant no. JPMJCR1654 (to M.M.).

## AUTHOR CONTRIBUTIONS

Conceptualization, H.I. and M.M.; Methodology, H.I., S.T., T.H., Y.K., C.O., S.T., S.I., A.M., K.S., and M.M.; Validation, H.I., T.H., K.T., and M.M.; Formal Analysis, H.I. and M.M.; Investigation, H.I., S. T., Y.K., and M.M.; Data Curation, H.I., T.H., and M.M.; Resources, H.I., C.O., S.T., S.I., A.M. K.S., and M.M.; Writing – Original Draft, H.I.; Writing – Review & Editing, T.H., A.M., K.S., K.T., and M.M.; Supervision, M.M.; Project Administration, M.M.; Funding Acquisition, H.I. and M.M.

## DATA AVAILABLITITY STATEMENT

Reagents, code and quantified data are available upon request.

## DECLARATOIN OF INTERESTS

The authors declare no competing interests.

## Supplemental Figures

**Figure S1:**
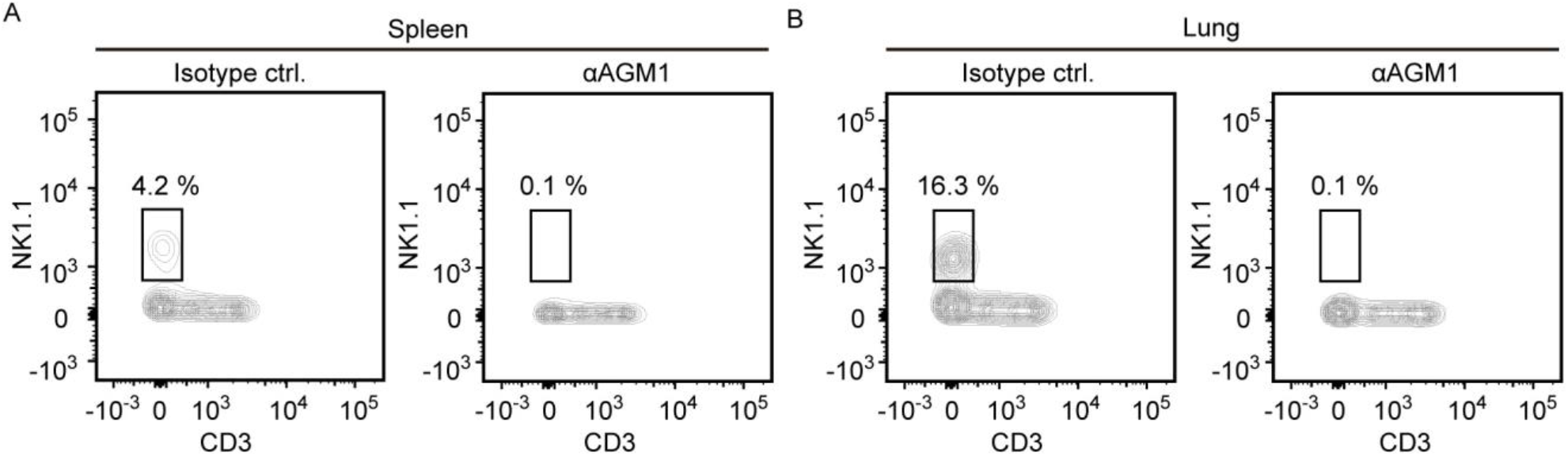
Depletion of NK cells by αAGM1. Flow cytometric analysis of the spleen (A) or lung (B) of mice treated with isotype control antibody or αAGM1. The numbers over the boxes indicate the percentage of NK1.1-positive cells among live single cells. Data are representative of 3 mice each.

**Figure S2:**
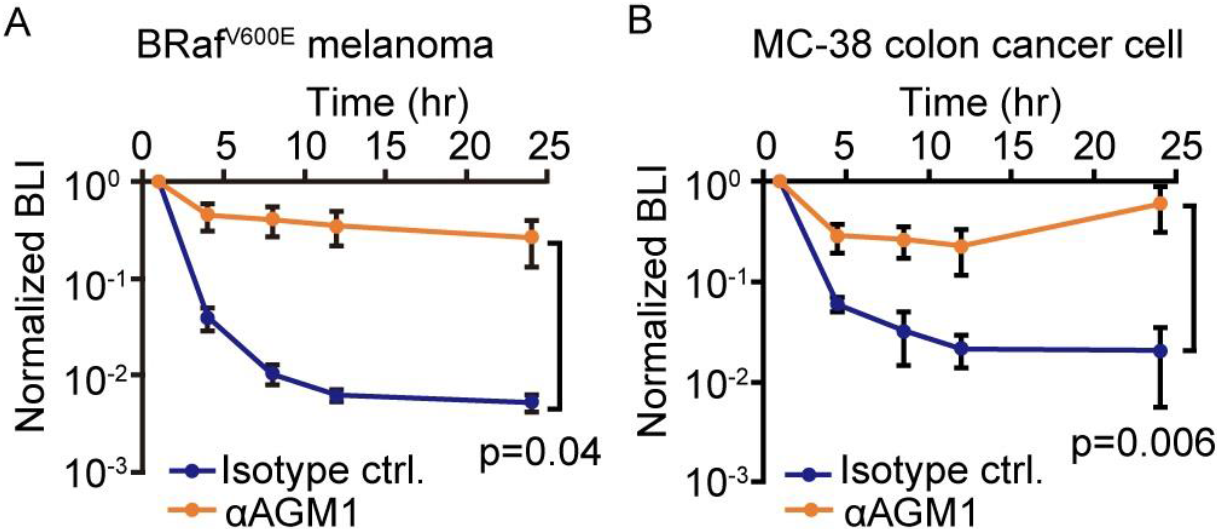
NK Cells Eliminate BRaf^V600E^ Melanoma and MC-38 Cells from the Lung within 24 hrs. BRafV600E melanoma (A) and MC-38 cells (B) expressing Akaluc were injected into the tail vein of mice treated with either control antibody or αAGM1. BLI was quantified at the indicated time and normalized to that at 1 hr after injection of the tumor cells. Data are representative of 3 independent experiments with 3–6 mice per group and are shown as means ± SD.

**Figure S3:**
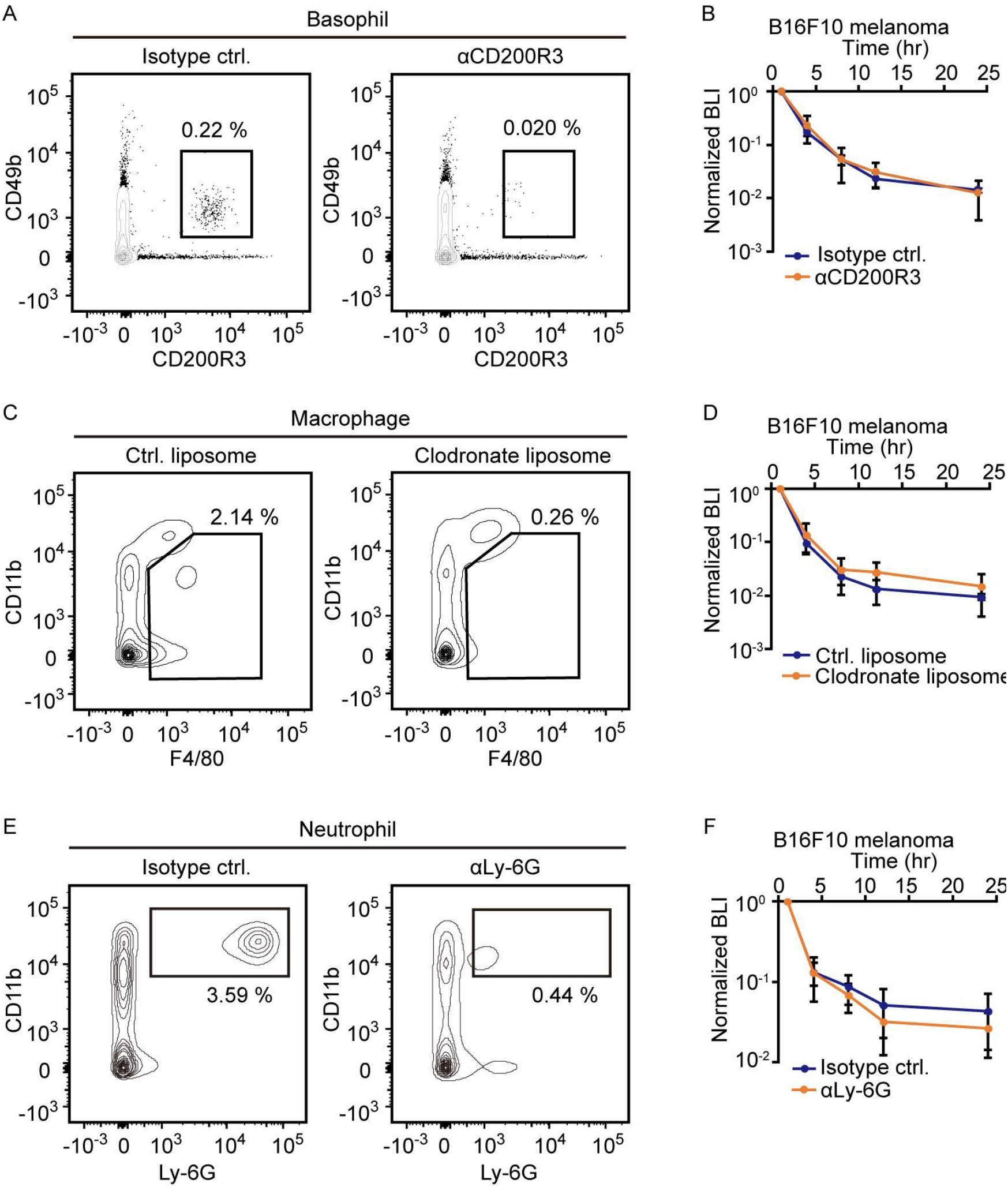
Basophils, Macrophages, and Neutrophils Do Not Contribute to Elimination of Metastatic Tumor Cells. Flow cytometric analysis of the spleens and bioluminescence imaging in mice depleted of basophils (a, b), macrophages (c, d), and neutrophils (e, f), by αCD200R3 antibody, clodronate liposome, and αLy-6G antibody, respectively. Data are representatives of three mice each for flow cytometric analysis and 2 independent experiments with 3 mice per group and are shown as means ± SD for bioluminescence imaging.

**Figure S4:**
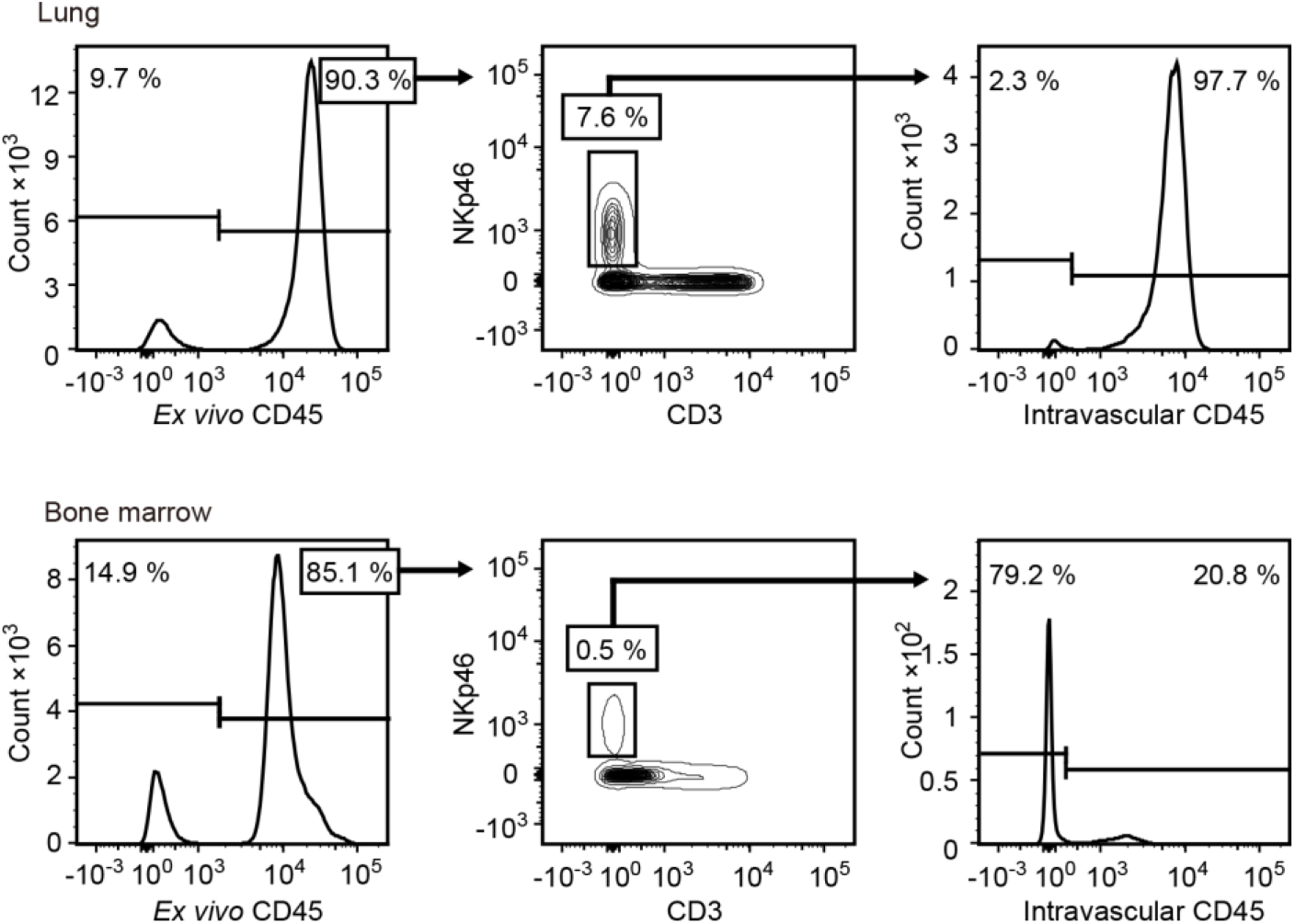
Intravascular Staining of NK Cells. Flow cytometric analysis of the lung (top) and bone marrow (bottom) of mice intravenously injected with αCD45 BV510 antibody and counterstained *ex vivo* with αCD45 FITC antibody. **Left**, Histogram of *ex vivo* CD45 expression on a live single cell gate. **Center**, Counter plots of CD3 and NKp46 expression on *ex vivo* CD45^+^ cells. **Right**, Histogram of intravenously injected CD45 antibody on CD3^−^ NKp46^+^ cells. Data are representative of 3 mice each.

**Figure S5:**
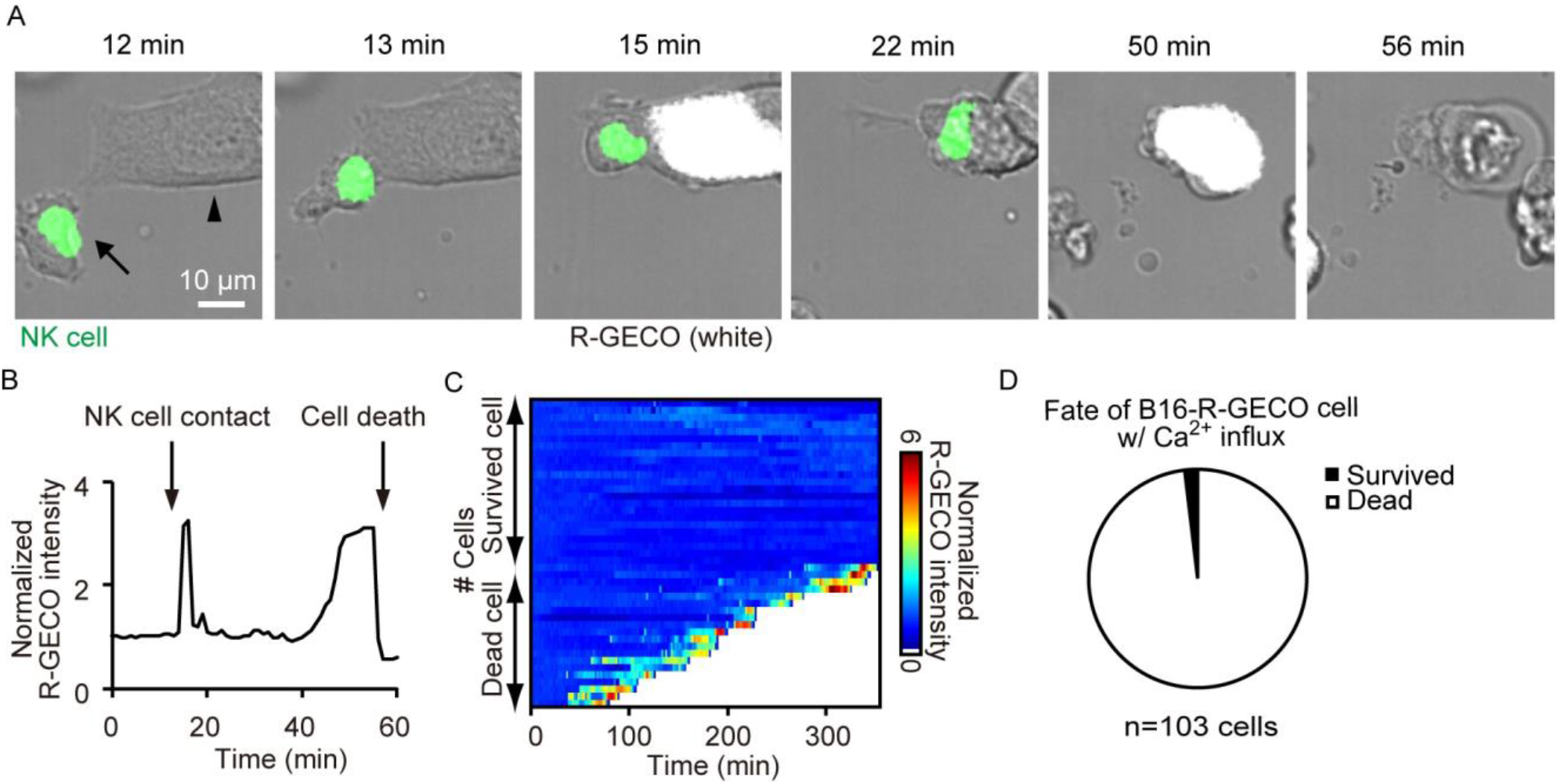
NK Cell-Induced Ca^2+^ Influx in B16F10 Cells *in vitro*. (A) A representative time-lapse image of the interaction between an NK cell (arrow) and a B16-R-GECO cell (arrowhead) *in vitro*. Shown here are merged images of differential interference contrast, YFP fluorescence (NK cell, green) and R-GECO1 fluorescence (B16-R-GECO cell, white). (B) Time course of R-GECO1 intensity. (C) R-GECO1 intensity in each cell is normalized to the intensity at the start of imaging and displayed as a heatmap. n=43 cells from 2 independent experiments. (D) Percentage of deceased tumor cells that exhibited Ca^2+^ influx after NK cell engagement. n=103 from 6 independent experiments.

**Figure S6:**
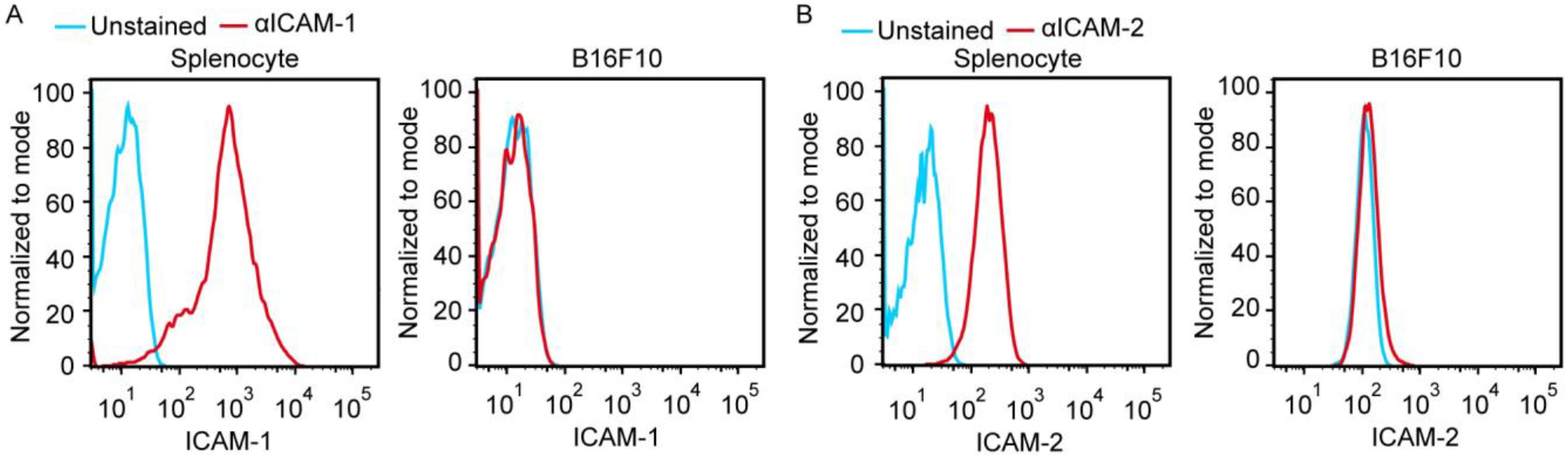
Absence of LFA-1 ligands on B16F10 cells. Flow cytometric analysis of the expression of ICAM-1 (A) and ICAM-2 (B) on the B16F10 cells.

**Figure S7:**
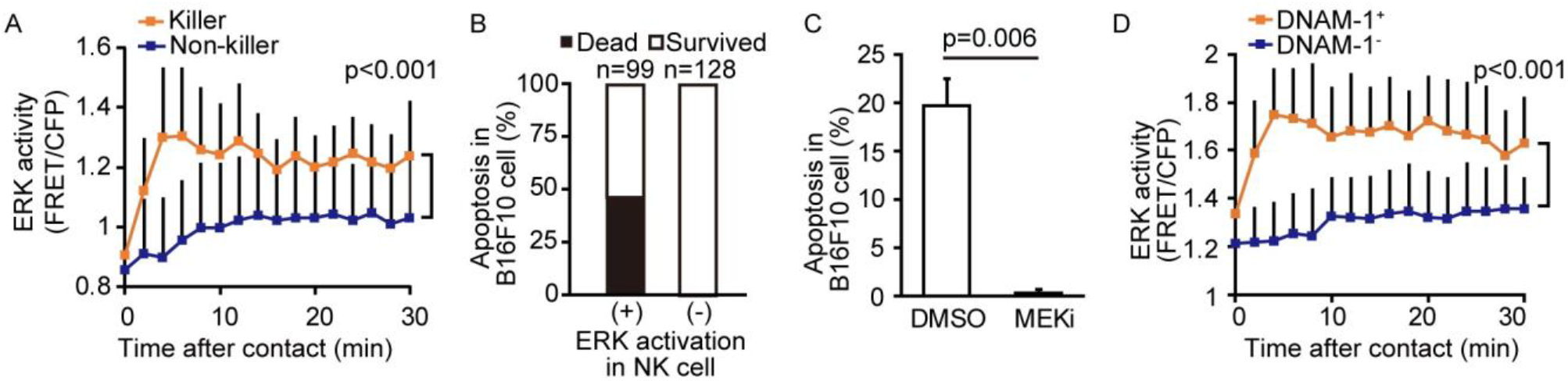
ERK Activation in the Killer NK Cells *in vitro*. (A) NK cells derived from hyBRET-ERK-NLS mice were cultured with B16-R-GECO and observed under an epifluorescence microscope. Quantification of the FRET/CFP ratio in the NK cells that induced apoptosis (killer cells) and those that failed to induce apoptosis (non-killer cells) in the target cells. Data were pooled from 6 independent experiments and are shown as means ± SD; n=43 cells for killer cells and n=73 cells for non-killer cells. (B) Induction of apoptosis in the target cells by NK cells with or without ERK activation. Data are from 6 independent experiments. (C) NK cells were cultured with B16-R-GECO cells in the presence or absence of MEKi. Percentages of target cell death are shown. Data are pooled from 3 independent experiments and represented as means ± SDs. (D) NK cells are sorted by the expression of DNAM-1. The DNAM-1^+^ or DNAM-1^−^ NK cells were cultured with B16-R-GECO cells. Data were pooled from 2 independent experiments and are represented as means ± SD; n=37 cells for DNAM-1^+^ cells and n=27 cells for DNAM-1^−^ NK cells.

**Figure S8:**
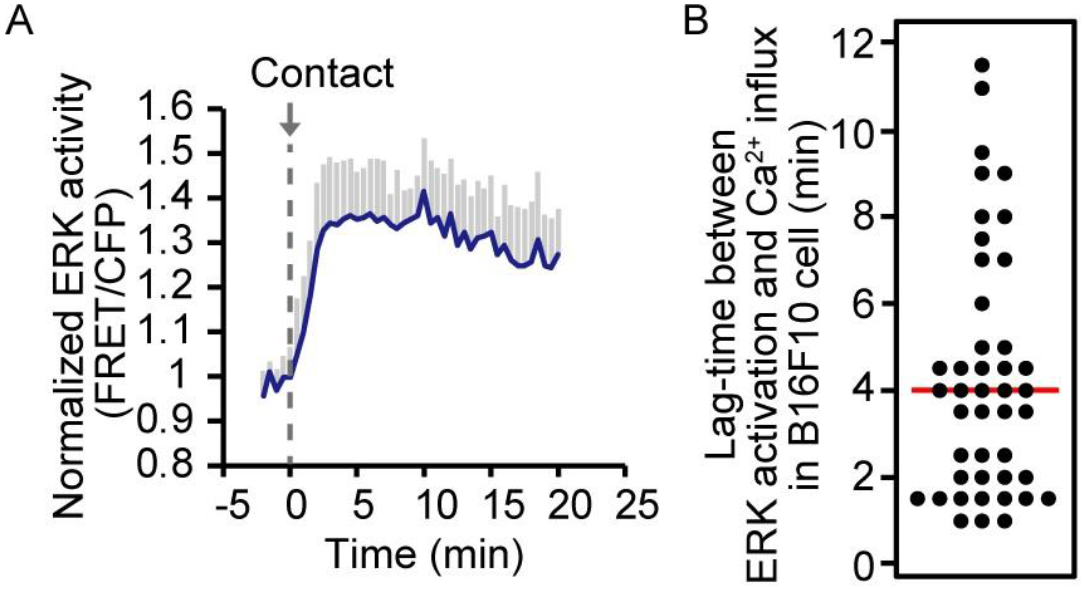
*In vivo* Dynamics of ERK Activity in NK Cells After Target Cell Contact. (A) Quantification of the FRET/CFP ratio in the NK cells that exhibited ERK activation after target cell contact. Data are pooled from 2 independent experiments and are represented as means ± SD, n=18 cells. (B) Time intervals between ERK activation in NK cells and Ca^2+^ flux in melanoma cells. Data were pooled from 3 independent experiments. A red line represents the mean. n=43 cells.

**Figure S9:**
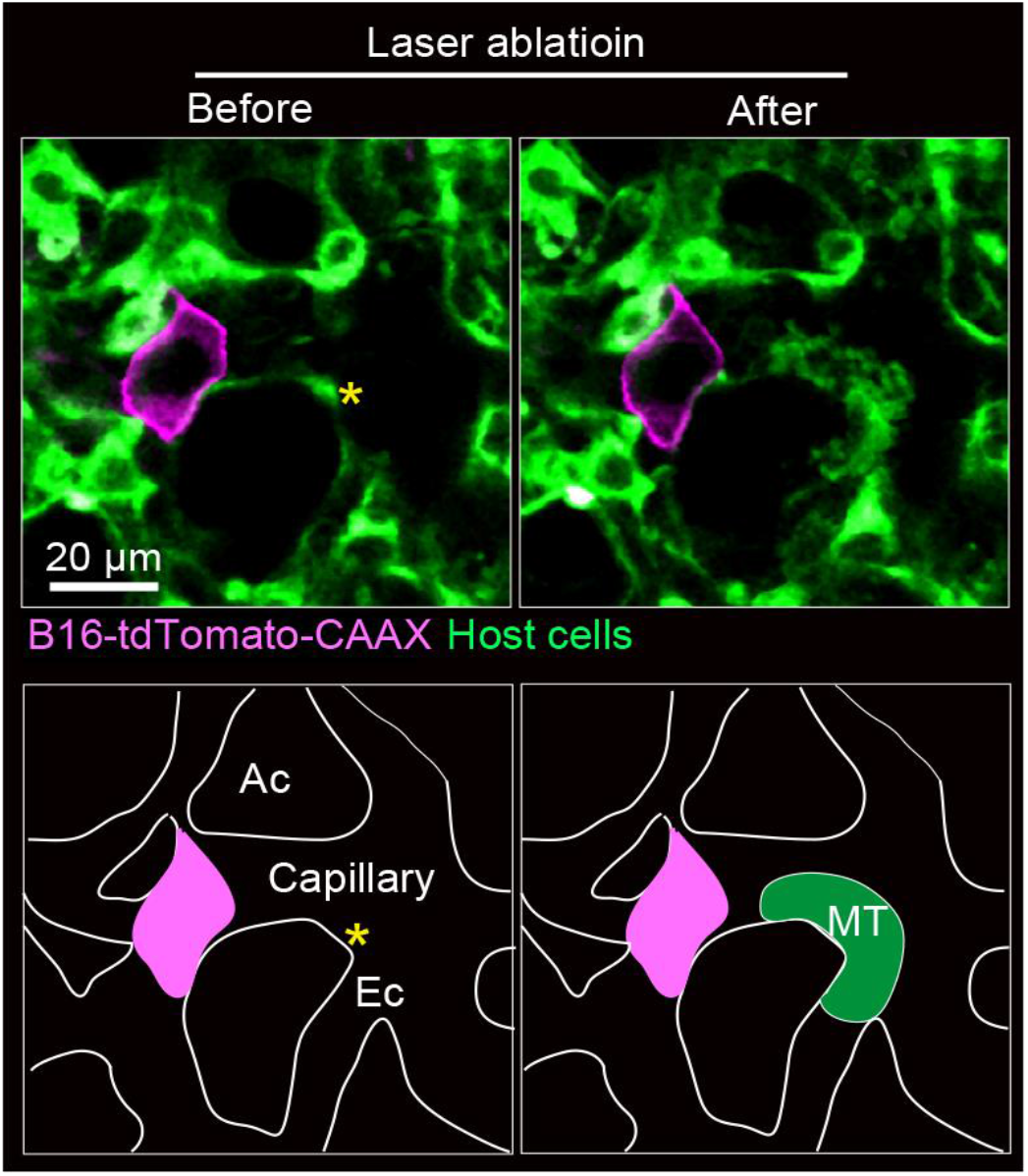
Lack of Micro-Thrombus Around the Disseminated Tumor Cells. In vivo imaging of pulmonary capillaries of a hyBRET-ERK-NES mouse, which was injected B16F10 cells expressing tdTomato-CAAX 4 hrs before imaging. The representative merged image of host cells (Green) and B16F10 cells (Magenta) is shown, with a schematic view of this region. Asterisks represent the laser ablated regions. Ac, alveolar cavity; Ec, endothelial cell. The image is representative of 2 independent experiments with 22 FOVs. See also Supplementary Video 5.

## Supplementary Table

Dynamics of NK cell killing of melanoma cells in the lung.

| Symbols | Parameters | Values | Units | References, equations, and comments |
| --- | --- | --- | --- | --- |
| **Histological and physiological parameters from published papers** |  |  |  |  |
| Bv | Blood volume | 1.7E-6 | m<sup>3</sup> | Table II(Davies and Morris, 1993) |
| Lv | Lung volume | 3.7E-07 | m<sup>3</sup> | Fig. 2A for week 8 mice(Gomes et al., 2019) |
| CO | Cardiac output | 2.0E-05 | m<sup>3</sup>/min | Abstract(Janssen et al., 2002) |
| Cp_l | Total lung capillary length | 1.1E+03 | m | Result section(Knust et al., 2009) |
| **Parameters determined experimentally** |  |  |  |  |
| FOV | Field of view | 2.5E+07 | m<sup>2</sup> | 0.5 x 0.5 mm |
| Plt_speed | Platelet speed | 0.057<br>0.95 | m/min<br>mm/sec | Determined as described previously(Sano et al., 2016). |
| Cp_r | Capillary radius | 3.4E-06 | m | Measured on the images of Fig. 2 |
| NK_d | NK cell diameter | 1.0E-5 | m | Measured on the images of Fig. 2 |
| NK_bl | NK cell count in blood | 5.1E+10 | cells/m<sup>3</sup> | Determined for C57BL/6 mice of 8–12 weeks old. |
| NK_FOV | NK cell number in a field of view | 16 | cells/FOV | Measured on the images of Fig. 3 |
| NK_speed | NK crawling speed on capillaries | 4.8E-06 | m/min | Determined with time-lapse images of Fig. 3 |
| NK_hit_obs | Observed NK cell hit probability | 8.0E-3 | cells/min | Determined with time-lapse images of Fig. 4 |
| NK_kill | NK cell killing probability | 0.5 |  | Determined with time-lapse images of Fig. 4 |
| MI_hl_BLI | Melanoma half-life based on BLI data | 146 | min | From bioluminescence images of Fig. 1B at 4 hrs. |
| **Calculated parameters** |  |  |  |  |
| Cp_fr | Capillary flow rate | 1.60E-12 | m<sup>3</sup>/min | Plt_speed* π*Cp_r<sup>2</sup> |
| NK_wbl | Whole blood NK cells | 8.7E+04 | cells | NK_bl*Bv |
| NK_density | NK cell density in lung | 6.4E+12 | cells/m<sup>3</sup> | NK_FOV/FOV/NK_d |
| NK_lung | Total NK cell in lung | 2.4E+06 | cells | NK_density*Lv |
| NK_in | NK cell influx to lung | 1.0E+06 | cells/min | CO*NK_bl |
| NK_out | NK cell decay constant | 0.42 | /min | NK_in/NK_lung |
| NK_hl | NK cell half-life in the lung | 1.6 | min | ln2/NK_out |
| NK_fr | NK cell flow rate per capillary | 0.11<br>6.6 | cells/min<br>cells/hr | NK_bl*Cp_fr |
| NK_dcp | NK cell density on capillary | 2.1E+03 | cells/m | NK_lung/Cp_l |
| NK_hit_cr | Crawling NK cell hit probability | 0.010<br>0.6 | cells/min<br>cells/hr | NK_dcp*NK_speed |
| MI_t_2P_obs | Melanoma decay constant calculated by 2P imaging | 4.0E-03 | /min | NK_kill*NK_hit_obs |
| MI_hl_2P_obs | Melanoma half-life based on 2P imaging data | 173 | min | ln2/MI_t_2P_obs |
| MI_t_2P_crsp | Melanoma decay constant based on crawling speed | 5.1E-03 | /min | NK_kill*NK_hit_cr |
| MI_hl_2P_crsp | Melanoma half-life based on crawling speed | 137 | min | ln2/MI_t_2P_crsp |

## Basic parameters

Macroscopic and histological data were based on previous papers (Gomes et al., 2019; Janssen et al., 2002; Knust et al., 2009). The speed of platelets in the lung capillaries (Plt_speed) was determined as described previously (Sano et al., 2016). The speed of platelets in the pulmonary capillaries was roughly one third of the speed of platelets in the arteriole of mouse bladder, 3.1 mm/sec (0.186 m/min) (Sano et al., 2016). The capillary radius, diameter of NK cells, and number of NK cells in a field of view were determined on at least 3 images. Plt_speed and Cp_r were used to calculate the capillary flow rate (Cp_fr). To determine the total number of NK cells in the blood, 50 μL of blood was collected from the right ventricle of 8–12 week-old C57BL/6 mice, lysed in ACK buffer (155 mM/L NH_4_Cl, 10 mM/L KHCO_3_, 0.1 mM/L EDTA), and analyzed by flow cytometry. The CD3^−^ NK1.1^+^ cells were counted as NK cells. This number of NK cells is two to three-fold larger than that reported previously by using C57BL/6J mice (Banh et al., 2012). The mean crawling speed on the endothelial cells is described in the text related to Fig. 3E. The probability of an NK cell hitting a tumor cell was determined by a MATLAB script (Main_191017.m). The probability of an NK cell killing a tumor cell was 0.5, based on the probability of induction of calcium influx in the target B16 melanoma cells (Fig. 4H).

## Total number of lung NK cells

The total number of NK cells residing in the lung (NK_lung) was estimated from the mean NK cell density (NK_density) and total lung volume (Lv). The diameter of an NK cell (NK_d) was used as the thickness of the image plane. The number of total NK cells, 2.4 million, is markedly larger than the previous values, which ranged from 0.2–1 million (Bi et al., 2017; Gregoire et al., 2007; Yan et al., 2014). In previous studies, the whole lungs were lysed to count the blood cell number. It is possible that the recovery rate might have been low due to insufficient tissue lysis. As described in the main text, we observed comparable numbers of tumor cells and NK cells in each FOV, when 1.5×10^6^ B16-SCAT cells were injected into NK-tdTomato mice, supporting the fidelity of the number of total NK cells determined in this study.

## Dynamics of NK cells

Most of the pulmonary NK cells are within the vasculature (Fig. S4), and the number of pulmonary NK cells overwhelms that of NK cells in the blood. Thus, the total number of lung NK cells can be used as the total number of NK cells in the lung vasculature. NK cell influx into the lung (NK_in) is obtained from cardiac output (CO) and NK cell count in the blood (NK_bl). If all NK cells stay in the lung with equal probability, the apparent transit time in the lung, or NK cell half-life in the lung, is calculated as 1.6 min from NK_in and NK_lung. This value is markedly smaller than the tracking duration period observed in Figs. 3 and 4, indicating that the major population of NK cells in the blood go through the lung without adhesion to the endothelial cells. By using the capillary flow rate (Cp_fr) and NK cell count (NK_bl), the NK cell flow rate per capillary (NK_fr) is determined as 0.11 cells/min. Meanwhile, from the total length of capillaries (Cp_l) and the number of NK cells (NK_lung), NK cell density on the capillary (NK_dcp) is determined as 2.1 cells/mm. From NK_dcp and the crawling speed of NK cells, the probability of a tumor cell being hit by crawling NK cells (NK_hit_cr) becomes 0.010 cells/min. This value is approximately one-tenth of the flow rate of NK cells (NK_fr).

## Dynamics of disseminated melanoma cells

The BLI signals from 1 to 12 hrs (Fig. 1B) were fitted with a built-in function of MATLAB (Fitting_1b.m) and obtained using the following equation:

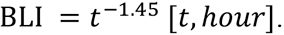

With this fitting, the decay rate decreases with time. Because we characterized the NK cell interaction with tumor cells between 4 to 8 hrs after tumor cell injection, we determined the half-life of B16F-Akaluc cells from 4 hrs (Ml_hl_BLI) and obtained 146 min. Meanwhile, from the probability of an NK cell hitting a tumor cell (NK_hit_obs) and the probability of an NK cell killing a tumor cell (NK_kill), the half-life of tumor cells (Ml_hl_2P_obs) becomes 173 min. If we adopt the probability of a tumor cell hit based on the crawling speed of NK cells, the expected half-life of tumor cells (Ml_hl_2P_crsp) becomes 137 min. Considering the precision of parameters obtained from *in vivo* imaging data, we believe that the half-life of melanoma cells estimated from the 2P microscopy reasonably matched the half-life of melanoma cells determined by BLI.

## Supplementary Video Legends

**Supplementary Video 1: Acute rejection of metastatic tumor cells by NK cells.**

The video corresponds to Fig. 1A. Merged images of the bright field and the bioluminescence images of mice intravenously injected with 5×10^5^ B16-Akaluc cells. Substrate was i.p. administered immediately after injection of tumor cells. Image acquisition was started at 5 min after tumor injection. Bioluminescence intensity is displayed in pseudo-color.

**Supplementary Video 2: Stall-crawl-jump movement of NK cells in the pulmonary capillary.**

The video corresponds to Fig. 2C. An example of an NK cell that moves within a pulmonary capillary in a stall-crawl-jump manner. A white arrow points to an NK cell showing a stall-crawl-jump movement.

**Supplementary Video 3: Induction of caspase 3 activation by crawling NK cells.**

The video corresponds to Fig. 4A. A time-lapse movie of the lung of an NK-tdTomato mouse after intravenous injection of B16-SCAT3 cells. The CFP/FRET ratio in B16-SCAT3 cells is shown in the IMD mode and an NK cell is shown in white.

**Supplementary Video 4: Induction of calcium influx by crawling NK cells.**

The video corresponds to Fig. 4E. Intravital imaging of the pulmonary capillary of NK-tdTomato mice after intravascular injection of B16-GCaMP cells. GCaMP6s intensity is displayed in pseudo-color. An NK cell is shown in white. A white arrow points to the melanoma cell that exhibits Ca^2+^ influx after contact with an NK cell (yellow arrowhead).

**Supplementary Video 5: Thrombus-formation by laser ablation around the disseminated tumor cells.**

The video corresponds to Fig. S9. Intravital imaging of the pulmonary capillary of hyBRET-ERK-NES mouse, which was injected B16F10 cells expressing tdTomato-CAAX 4 hrs before imaging. Host cells and tumor cells are shown in green and magenta, respectively.

